# Parallel phenotypes underpinned by different genes in the visual system of two trans-isthmian coral reef fish species pairs

**DOI:** 10.64898/2026.06.08.730927

**Authors:** Michele E. R. Pierotti, Eirlys E. Tysall, Marc P. Hoeppner, Christopher Haak, Ryan A. Vandermeulen, Henry Goehlich, Ellis R. Loew, D. Ross Robertson, Karen L. Carleton, W. Owen McMillan, Andrea Manica

**Author notes:** Corresponding Authors: Michele E. R. Pierotti, Eirlys E. Tysall. These authors contributed equally to this work (Michele E. R. Pierotti, Eirlys E. Tysall). Joint senior authors (W. Owen McMillan, Andrea Manica).

## Abstract

Understanding the genetic basis of adaptation in natural populations to changing environmental conditions is challenging. The relatively simple genotype-to-phenotype relationship between opsin genes and visual pigments offers a particularly informative model to explore the molecular mechanisms underpinning adaptive phenotypic change in natural populations. Here we leveraged the natural experiment provided by the tectonic closure of the Central American seaway that created allopatric, sister taxa in multiple, independent lineages of marine organisms and exposed them to distinct underwater light environments: either the more turbid Tropical Eastern Pacific (TEP), or the spectrally broader Caribbean Sea. Using two species pairs of planktivorous teleosts, the *Azurina multilineata*/*A. atrilobata* damselfish and the *Cephalopholis (Paranthias) furcifer*/*C. colonus* groupers, we explore to what extent visual sensitivity converged to similar adaptations in response to similar foraging strategies and shared underwater light in each marine basin. We found that the compression of the underwater light field towards the central portion of the spectrum from Caribbean to TEP waters is reflected in similar shifts towards the centre of the light spectrum in overall single and double cone sensitivities in both families. Both TEP species have single (short-wavelength) cone sensitivities shifted to longer wavelengths and double (long-wavelength) cone sensitivities shifted to shorter wavelengths, compared to their Caribbean counterparts. These parallel shifts in visual sensitivities observed in response to shared underwater light environments are accomplished by different underlying opsin gene toolsets in the two lineages. Similarly, expression changes in pathways associated with the visual system revealed limited parallelism at the molecular level.

## Introduction

Understanding adaptation and functional divergence in response to changing environmental conditions is a major goal of evolutionary ecology (Levins 1968, Bomblies and Peichel 2022). However, discerning which traits divergent selection acts upon, the molecular mechanisms underlying phenotypic change and the ecological conditions driving such changes can be challenging, especially in natural populations (Kawecki and Ebert 2004, Savolainen et al. 2013, Wadgymar et al. 2017).

Vision is arguably one of the best understood systems in terms of the molecular mechanisms that underlie phenotypic change, providing an ideal target to study natural selection. In vertebrates, visual opsins, through their interactions with the light-sensitive pigment retinal, modulate the selective absorbance of light reaching the retina, and thus play a central role in modulating dim-light and colour vision. As the sensitivity spectrum of the photoreceptor is largely determined by the amino acids of the opsin, there is a direct link between genotype (the genetic sequence of opsin) and the phenotype (the sensitivity of individual photoreceptor cells to certain wavelengths). This one-to-one map and the extensive body of mutagenesis experiments detailing the effects of single amino acid substitutions on spectral sensitivity (Yokoyama and Yokoyama 1996, Yokoyama 2000, 2008, Yokoyama et al. 2008), make colour vision an ideal system to study the effects of divergent selection on a well-defined phenotypic trait with a simple genetic basis, in natural populations (Terai et al. 2006, Tezuka et al. 2014, Kawamura et al. 2016). Beyond the amino acid sequence and repertoire of opsin genes, changes in the relative expression of each opsin class provide an additional mechanism of tuning visual sensitivity in response to environment change (Carleton and Kocher 2001, Hofmann et al. 2009, Rennison et al. 2016).

The emergence of similar traits in multiple lineages experiencing similar environments has long interested evolutionary biologists as potential examples of adaptation in response to a common selective pressure (e.g. Darwin 1959, Schluter and Nagel 1995, Gould 2002, Rosenblum et al. 2014, Stuart et al. 2017, Bolnick et al. 2018, Greenway et al. 2020). Comparing the genomes of three-spine stickleback (*Gasterosteus aculeatus*) that repeatedly invaded either clear, oligotrophic or tannin-rich blackwater lake populations, Marques et al. (2017) found that the same red-shifting alleles in the blue-sensitive opsin *SWS2A* facilitated the multiple, independent invasions of blackwaters. This result was further supported by a powerful, 19 years-long selection experiment where sticklebacks were transplanted from blackwater to clearwater habitats. Even more remarkable is that the same amino acid replacements at the two tuning sites conferred blue-or red-shift in the stickleback *SWS2A* replicate, marking a striking example of parallelism (Bolnick et al. 2018). Variation at the same site between ancient paralogs, a blue-shifted *SWS2B* and a red-shifted *SWS2A*, is present in other teleosts (Yokoyama et al. 2007, Cortesi et al. 2015). By contrast, in a study of colonization of the same oligotrophic crater lake from turbid rivers and shallow lakes, Härer et al. (2018) found no evidence of recurring recruitment of the same opsins and tuning sites in the seven independent invasions of clear waters by Nicaraguan cichlid fish species. Gene expression patterns across photic environments also differed between species though they often resulted in similar (parallel) phenotypic shifts in overall visual sensitivity.

The emergence of the Isthmus of Panama leading to the closure of the Central American Seaway around 3 MYA (Coates et al. 2005, O’Dea et al. 2016) provides a natural experiment to study replicated adaptation to changing underwater light environments in taxonomically diverse assemblages i.e. moving beyond comparisons between populations or closely related species (e.g. Marques et al. 2017, Härer et al. 2018). Whilst the isthmus created a permanent bridge for terrestrial species, setting in motion the Great American Interchange (Stehli and Webb 2013), it also gave rise to a complete physical barrier to gene flow for marine species with populations previously connected across either side of the Americas. This not only acted as a major vicariant event for marine taxa but progressively determined changes in water circulation, radically modifying the oceanographic characteristics of the separated water bodies, in particular, temperature, salinity and nutrients. The Caribbean today is a warm, oligotrophic and relatively stable sea, where a broad light spectrum penetrates to substantial depths. By contrast, the closure of the isthmus brought the Tropical Eastern Pacific (TEP) under the influence of seasonal upwellings, intense primary productivity and high concentrations of dissolved organic matter (D’Croz and O’Dea 2007, 2009). These environmental differences result in very different light environments in the two oceans. Characteristics of the spectrum of underwater light (Munz and McFarland 1977, Shand et al. 2008, Schweikert and Grace 2018) and its attenuation by absorption and scatter (Johansen and Jones 2013, Ehlman et al. 2015, Chang and Yan 2019) are known to exert strong selective pressures on aquatic visual systems. The short, yet recent time frame of the closure, and the global scale of its effects, created entirely new biogeographical provinces. This event provides us today with an abundance of study systems on which to examine the effects of vicariance and divergent ecological selection in marine species. Here, we take advantage of this natural experiment and examine the divergence in the light environment and visual system of pairs of allopatric sister taxa. The so-called trans-isthmian geminates (Jordan 1908) live in either a stable, oligotrophic and blue-shifted sea, the Caribbean, or in the nutrient-rich, more turbid Tropical East Pacific. We expect these sister species to have been separated for at least the ∼3 million years following the closure of the isthmus, though there is potential for divergence to pre-date the final closure. We focus on two geminates belonging to very distant lineages of teleosts: the planktivorous damselfish (family Pomacentridae) *Azurina multilineata* in the Caribbean (CAR) and *Azurina atrilobata* in the Tropical Eastern Pacific (TEP), and the groupers (family Serranidae) *Cephalopholis (Paranthias) furcifer* (CAR) and *Cephalopholis (Paranthias) colonus* (TEP) (Fig. 1A). We also include their closest extant relatives, namely, *Azurina cyanea* and *Cephalopholis fulva*, both co-occurring in the Caribbean and, as *C. furcifer* and *A. multilineata*, more widely, in the Tropical Western Atlantic. Whilst the damselfish geminates belong to a whole lineage of specialised planktivores, the Chrominae (Frédérich et al. 2016), the grouper geminates are unusual in having recently abandoned the crepuscular ambush predator niche typical of the genus *Cephalopholis* and most mid-sized groupers (including *Cephalopholis fulva*). Instead, they invaded a new habitat and food source, becoming diurnal open-water planktivores, foraging on the same reefs and on overlapping diet items (Randall 1967; Fig. S1) with the *Azurina* geminates. We are thus able to compare how the visual system of these two distant geminate pairs, with very similar ecologies, but visual systems historically shaped by very different evolutionary forces and constraints, have responded to the shared selective pressures imposed by the same shift in photic environment between the two ocean basins.

**Figure 1.**
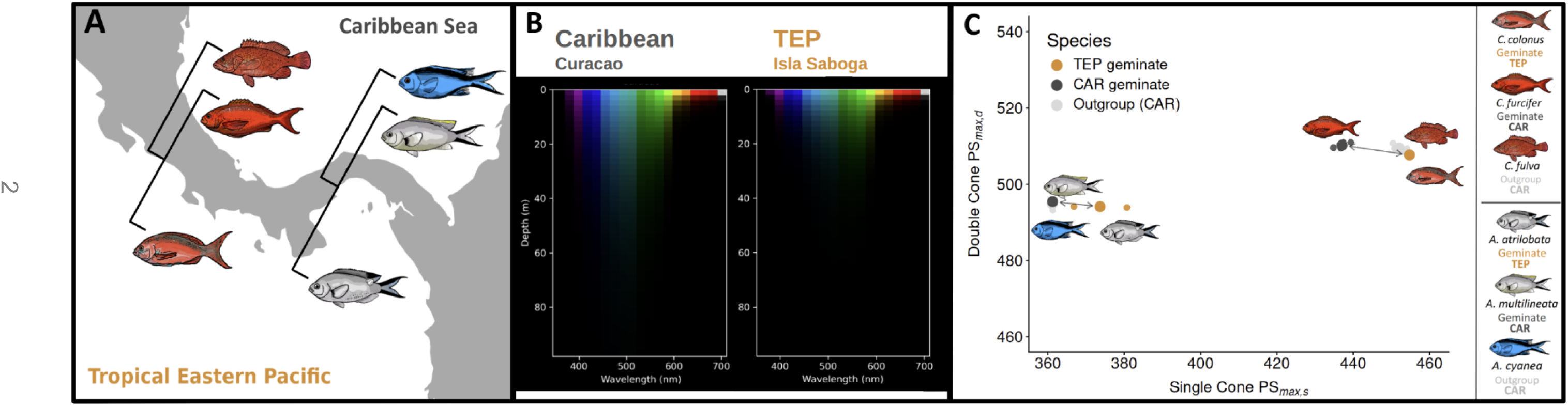
A) The Isthmus of Panama and study species. Each pair of sister species (‘geminates’) consists of one member in the tropical Eastern Pacific and one in the tropical Western Atlantic. In addition, the closest extant relative of each geminate pair is included in the study; **B)** Approximation of the underwater light field at depth based on the remote sensing estimated irradiance profiles taken on February 2025 at the collection sites: Curacao (CAR) (left) and Isla Saboga (TEP) (right); **C)** The combined effect of cone spectral shifts and changes in cone gene expression as expressed by the maximal sensitivity index PS*_max_* for single versus double cones in each geminate species pair and their outgroups. Arrows between the tropical Eastern Pacific (TEP) and Caribbean Sea (CAR) geminate species indicate the parallel shift in maximal sensitivity in both damselfish and grouper pairs despite large differences in the underlying opsin repertoire.

To understand the response of the visual system to the contrasting optical properties of the CAR and the TEP, we first characterised underwater light environments directly using in situ irradiance measurements and by modelling remote-sensing reflectances across multiple sites in both CAR and TEP during the dry and wet seasons, over a ten-year period. We then quantified phenotypic adaptation by measuring the spectral absorbance of individual photoreceptors in the retina with micro-spectrophotometry. We studied the genetic changes underpinning such phenotypic changes by looking at amino acid changes in the coding sequence of visual opsins, taking advantage of newly assembled, high quality reference genomes for each species. Finally, we used transcriptomes from four individuals of each species to characterise expression differences that might have complemented such sequence changes in the opsins, as well as differences in pathways that are associated with vision. When combined, these lines of evidence provide a comprehensive picture of repeated adaptive divergence of the visual system in two phylogenetically distant pairs of sister species (the geminates), under the selective pressure of a clear, blue water environment or that of nutrient-rich and turbid, green waters.

## Results

### Light spectral field of the two basins

We measured downwelling, upwelling and sidewelling irradiances *in situ* at four depths, at the sites (Isla Saboga, TEP; Curacao, CAR) where specimens were collected for retina micro-spectrophotometry and genomic analyses (Fig. S2, Table S1). In addition, to identify basin-wide trends in spectral light field distribution, we obtained remote sensing reflectance data over multiple years and at various localities across CAR and TEP (Fig. S3), then used those to generate estimates of downwelling irradiances at any depth.

Both *in situ* measurements and remote sensing-based estimates show lower irradiance levels in the more nutrient-rich TEP than the oligotrophic CAR (Figs. 1B, S2A, S5, S6A; Table S1), and a substantial narrowing of the light field and red-shift in the wavelengths at which photons are most abundant, in TEP waters, at all depths (Figs. S2B, S2C, S4, S5, S6B). The in-situ measurements suggested a narrower spectral bandwidth (Δλ) in the TEP, with a loss of the deeper blue and green-yellow wavebands, as a result of wavelength-dependent absorption and scatter (Fig. S2C). This pattern was particularly evident for the horizontal (i.e., sidewelling) and downwelling light, the most relevant for an open-water, plankton feeding fish (Fig. S2; Table S1). Furthermore, the remote sensing irradiances indicated that lower underwater light levels and a red shift of the spectral field in the TEP compared to CAR conditions are characteristic features found at all measured stations across the two basins, over a time span of ten years, during or outside Panama’s upwelling season (Fig. S5), and therefore represent a regional pattern.

To facilitate comparisons between sites and seasons, we used the Apparent Visible Wavelength (AVW) (Vandermeulen et al. 2020): this is the weighted harmonic mean across visible wavelengths, summarising spectral shape with a single, intuitive value that expresses the dominant colour of downwelling light. Stations in both CAR and TEP exhibited higher (i.e. redshifted) AVW values in February, when local upwellings affect underwater light, than in September, when such climatic conditions are absent (Fig. S6A). Notably, TEP stations had significantly higher AVW values than CAR stations (red-shifted water colour index) in both seasons, over the 10-year timeframe, and at all depths evaluated (Fig. S6B).

Variation in average visible wavelength (AVW) at intermediate (30 m) depth was examined using a linear mixed-effects model with region (CAR vs TEP), season (dry vs wet), and their interaction as fixed effects and site as a random intercept. We found that the AVW differed significantly between regions (*F*₁,₁₀.₀ = 31.0, *p* = 0.0002) and between seasons (*F*₁,₂₁₉.₁ = 96.9, *p* < 0.0001), with the effect of season depending on region (*F*₁,₂₁₉.₁ = 72.4, *p* < 0.0001). In the dry season, the TEP had substantially higher (i.e. long wavelength-shifted) AVW than the CAR (TEP: 522 nm, CAR: 483 nm; difference = 39.2 nm, 95% CI: 27.3–51.2, *p* < 0.0001), and this difference between the two oceanic basins persisted into the wet season but was approximately half the magnitude (TEP: 501 nm, CAR: 482 nm; difference = 19.9 nm, 95% CI: 7.9–31.8, *p* = 0.004). The AVW in the CAR showed no significant changes between seasons (difference = 1.5 nm, 95% CI: −1.6–4.6 nm, *p* = 0.336), whereas it declined substantially in the TEP, from dry to wet season (difference = 20.9 nm, 95% CI: 17.7–24.1, *p* < 0.0001). Site accounted for 51.7% of the total variance in AVW (random intercept SD = 8.98 nm, bootstrap 95% CI: 5.10–13.01 nm; residual SD = 8.67 nm, bootstrap 95% CI: 7.86–9.50 nm; *n* = 233 observations from 12 sites). In conclusion, despite spatial heterogeneity in optical conditions among locations, when averaging across sites, the TEP is about 20nm ‘greener’ than the CAR during the wet season, and up to 40nm ‘greener’ than the CAR during the dry (upwelling) season, with the latter exhibiting only limited fluctuations in underwater spectral light across seasons.

Overall, these data show that differences in underwater light field between the two oceanic basins, and thus the potential for divergent selection on fish visual systems, are large, consistent across the two regions and across years, and they are not restricted to a particular season.

### Visual opsin repertoire

As expected, the visual opsin repertoire of the three damselfish species (family *Pomacentridae*) and the three grouper species (family *Serranidae*) were markedly different. In line with previous findings (Hofmann et al. 2012, Stieb et al. 2016), we recovered sequences for five cone opsins (*SWS1*, *SWS2B*, *RH2A*, *RH2B*, *LWS*), in addition to the rod opsin *RH1,* in the damselfish species. In the three groupers, we identified eight cone opsins: *SWS1*, *SWS2B*, *SWS2A1*, *SWS2A2, RH2B*, *RH2A1*, *RH2A2* and *LWS.* This is consistent with the rich cone class repertoire found in other grouper species: *Epinephelus moara* carries seven fully coding cone opsin genes (Musilova and Cortesi 2023) while *Epinephelus bruneus* has nine (Matsumoto and Ishibashi 2015). The identity of all recovered opsin sequences from all six species was confirmed using maximum likelihood trees including opsins from a range of teleosts (Figs. S7-S11; Table S2).

### Phenotypic changes in visual opsin sensitivity

In both geminate pairs, the peak sensitivity of individual photoreceptor cells, as directly measured through micro-spectrophotometry (MSP), was found to be shifted towards the centre of the spectrum in the TEP member when compared with its Caribbean counterpart (Fig. 2A, 2B; Table S3), despite the different underlying opsin gene repertoire between damselfish and groupers. This shift was particularly evident in the shorter-wavelength pigments (Fig. 2A, 2B; Table S3) where we expect the impact of the light environment on visual systems to be most prominent as a result of absorption and scatter. A more limited effect could also be observed in the long-wavelength opsins, with a clear blue-shift in *RH2A* in the damselfish (Fig. 2A) and *LWS* in the groupers (Fig. 2B).

**Figure 2.**
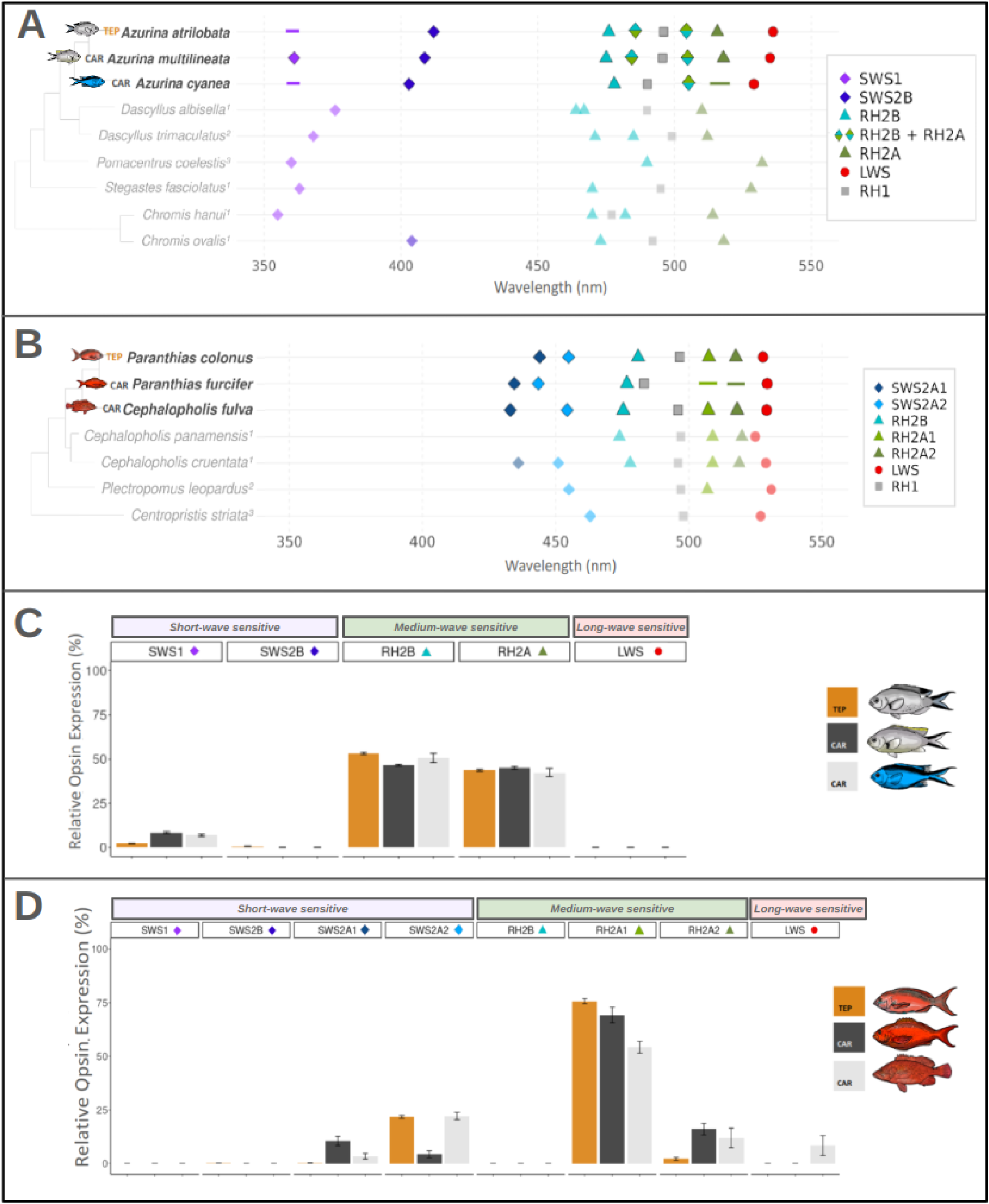
Wavelength of photoreceptor maximum absorbance (λ_max_) measured *in situ* with micro-spectrophotometry (MSP) and proposed relationship with expressed opsin genes, in **A**. the damselfish geminate pair and its outgroup, and **B**. in the grouper geminate pair and its outgroup. Cone class representation and associated maximum absorbance are placed in the context of available λ_max_ information (respectively, in A., from other members of the family Pomacentridae: ^1^Losey et al. (2003), ^2^Hawryshyn et al. (2003), ^3^McFarland and Loew (1994); in B., from other members of the family Serranidae: ^1^Pierotti unpublished data, ^2^Marshall et al. (2019), ^3^Singarajah and Hárosi (1992)). Instances where MSP measurements were not possible, but presence and expression of the opsin gene have been confirmed, are indicated with a line. **C, D)** Relative cone opsin expression as a percentage of total cone opsin expression, based on Transcripts Per Million (TPM) values, in the damselfish (C) and grouper (D) species. Error bars denote +/-standard error.

Based on cone spectral sensitivity, we identified up to eight spectral classes in the damselfish geminate pair and six in the grouper pair (Fig. 2A, 2B; Table S3). The mean and standard deviation of the wavelength of maximum absorbance λ_max_ and their inferred opsin gene class are presented in Supplementary Table S3. In the damselfish pair, we found the mean peak absorbance of the shorter-wavelength single cone pigment (putatively *SWS2B),* at a longer wavelength (λ_max_ = 412nm) in the TEP geminate than in both the Caribbean geminate (λ_max_ = 408.5 nm) and the Caribbean outgroup species (λ_max_ = 403nm) (Fig. 2A; Table S3).

Similarly, in the groupers, the mean peak absorbances of the two short-wavelength single cone pigments (putatively *SWS2A1* and *SWS2A2)* were both considerably blue-shifted between the TEP and Caribbean geminates. In the TEP species *C. colonus*, *SWS2A1* is shifted 10nm (λ_max_ = 444nm) to longer wavelengths compared to the Caribbean *C. furcifer* (λ_max_ = 434nm); the putative *SWS2A2* is red-shifted by about 11nm in *C. colonus* (λ_max_ = 454.9nm), as compared to *C. furcifer* (λ_max_ = 443.5nm) (Fig. 2B; Table S3).

At the other end of the spectrum, we found an opposite pattern in both geminates. The damselfish double cone green pigment (putative *RH2A*) peaked at slightly shorter wavelengths in the TEP geminate (λ_max_ = 515.7nm), compared with its Caribbean counterpart (λ_max_ = 517.9nm) (Fig. 2A; Table S4). In the grouper pair, the putative *LWS* carried in the double cones was slightly shorter in the TEP geminate (λ_max_ = 527.9nm) than in the Caribbean geminate (λ_max_ = 529.5) (Fig. 2B; Table S3).

Interestingly, in the damselfish geminates, two additional spectral classes were found in double cones, likely resulting from coexpression of *RH2A* and *RH2B* in two distinct proportions. First, a ‘cyan’ cone class was found in *A. multilineata* (λ_max_ = 484nm) and in *A. atrilobata* (λ_max_ = 486nm), but not in the outgroup *A. cyanea*. A second spectral group was identified with average λ_max_ = ∼507nm in all three species (Fig. 2A; Table S3). Thorough inspection of each species’ long-read genomic assembly did not find any evidence of additional *RH2* paralogs, supporting a co-expression scenario. Fitting experimental absorbance curves with combinations of putative pure *RH2A* and pure *RH2B* at the observed average λ_max_ indicates that two alternative co-expression ratios (1/3*RH2A* +2/3 *RH2B*; 2/3*RH2A* +1/3 *RH2B*), similar across species, can explain the two additional spectral classes.

### Sequence changes underpinning phenotypic adaptation

By analysing the coding regions of the opsin genes (as reconstructed from the *de novo* genome assemblies together with the *de novo* transcriptomes), we were able to identify a number of amino acid changes that underpin the phenotypic responses described above (Tables S4-S6). Whilst the phenotypic changes show parallelism, the amino acid changes involved differed between the two families. In the damselfish, the most prominent effects came from two tuning-site substitutions in *RH2B* (*multilineata* L207M *atrilobata*, S124A; Table S5) giving a 5nm red-shift; a mutation (F43L) giving a red shift of 3nm in the violet opsin *SWS2B;* a 5nm red-shift caused by two mutations in RH2B (L207M, S124A), and a substantial blue-shift of 16nm from a substitution in *LWS (multilineata* T282A *atrilobata)*. In the groupers, the largest shift was observed in the rod opsin *RH1*, where the combined effect of two substitutions (*colonus* A292S *furcifer*, S299A; Table S6) results in a 10nm shift in peak absorbance, consistent with the general pattern of longer wavelength rod sensitivity in the TEP. A further substitution in blue-sensitive opsin *SWS2A1* (*colonus* M116T *furcifer*) was present at a known tuning site, indicating a potential shift, though its magnitude is not known.

### Visual opsins expression

Complementary to the changes in opsin sequence, shifts in relative expression of each opsin class can lead to large spectral shifts, allowing the tuning of the visual system to the environment. We found consistent patterns among the two families of shifting expression indicative of adaptation to a narrower spectrum of light, with the TEP member of both geminate contrasts over-expressing opsins in the central region of the spectrum compared to their CAR twin. Comparable shifts in expression can be seen within the short-wave and medium-wave opsin complements available in both geminate comparisons (Fig. 2C, 2D).

In the damselfish, we found overall cone opsin expression to be *RH2*-dominated (Fig. 2C). Within this, we found varying ratios of expression between the longer (*RH2A*) and shorter (*RH2B*) wave sensitive pigments (Figs. 2C, S12B); whilst the CAR geminate *A. multilineata* expressed similar levels of *RH2B* and *RH2A*, the TEP geminate *A. atrilobata* expressed significantly more *RH2B* than *RH2A* in its cones. We recovered a similar pattern of narrowing expression within the short-wavelength region. Both CAR species had UV-sensitive *SWS1* overwhelmingly expressed in their single cones with only minimal expression of *SWS2B*. The TEP geminate on the other hand, had much lower expression of *SWS1* replaced in their single cones by *SWS2B*, suggesting that, as observed with the absorbance shifts, the TEP geminate tends to privilege opsin expression in a narrower, central region of the visual spectrum (Figs. 2C, S12A).

As observed in the damselfish, the groupers also had overall *RH2*-dominated cone opsin expression within which a broader palette was utilised by the CAR geminate, which expressed both *RH2A1* and *RH2A2,* while the TEP geminate’s expression was dominated by (shorter) *RH2A1* with little contribution from the more red-shifted *RH2A2* (Figs. 2D, S13B). At the shorter end of the spectrum the reverse pattern applied; within the single cones the CAR geminate expressed almost exclusively (shorter-wavelength) *SWS2A1* while the (longer wavelength) *SWS2A2* opsin was prevalent in the TEP geminate (Figs. 2D, S13A). Interestingly, whilst both grouper geminates possess an intact *SWS1* gene carrying the UV-tuning substitution (F86) (Cowing et al. 2002), we did not find any *SWS1* expression in either species. This absence is despite their typically planktivorous habits and seemingly overlapping diet with the damselfish; Caribbean grouper *C. furcifer* (no expression of the UV-sensitive *SWS1*) and Caribbean damselfish *A. multilineata* (UV-sensitive *SWS1* expressed) are known to feed largely on the same zooplankton species and within the same prey size ranges (Randall 1967; Fig. S1).

As expected, dim-light opsin *RH1* expression was highest in the damselfish outgroup *A. cyanea*, which ventures deeper in the water column; the geminates however showed very similar levels of expression (Fig. S14A). In the groupers, *RH1* levels reflected the overall light intensity in the habitats occupied by the three species, with higher expression in the TEP grouper geminate, subjected to lower light levels than its counterpart in the CAR, as well as higher expression in the demersal *Cephalopholis fulva* (Fig. S14B), consistent with its microhabitat and crepuscular habits.

### Predicted maximal sensitivity

The predicted maximal sensitivity (PS_max_), measuring the combined effects of a pigment’s wavelength of maximum sensitivity and its relative expression, was calculated for single and double cones, in both geminate pairs (Carleton et al 2008). Despite large differences in the underlying opsin repertoire between damselfish and groupers, we found parallel shifts in sensitivity between the TEP and the Caribbean member of each pair (Fig. 1C). In the single cones, typically carrying opsins on the shorter end of the spectrum, sensitivity in both damselfish and grouper pairs is shifted to longer wavelengths when comparing Caribbean to TEP geminate species. This shift tracks the loss of the shortest wavelengths in the turbid TEP (Figs. 1B, S2, S4), observed at all stations. At long wavelengths, spectral differences between TEP and CAR are less pronounced. Sensitivity in double cones, expressing the middle and longer wavelength opsins, shows limited but concordant shifts to shorter wavelengths in both damselfish and groupers.

When combined, these changes lead to an overall narrowing of the spectral sensitivity range in the TEP geminates, particularly at the shortest wavelengths, matching the shifting of the dominant component of the underwater light field in the TEP as compared with the broader spectrum of light available in the Caribbean.

The damselfish outgroup *A. cyanea* and the geminate *A. multilineata* are both planktivores, often foraging together on the same reefs in the Caribbean: as expected, we found their visual sensitivity to be very similar (Fig. 1C). Consistent with an ecological adaptive basis for sensitivity, a very different pattern from the damselfish outgroup is observed in the grouper geminates, both planktivores, and their CAR outgroup, *C. fulva*, an epibenthic predator. While the grouper planktivore geminates exhibit shifts in their visual system from CAR to TEP similar to those observed in the planktivore damselfish geminate pair, the grouper (CAR) outgroup showed a distinct, long-shifted visual system, reflecting the visual requirements of a bottom dwelling ambush predator (Fig. 1C).

### Transcriptome wide differential expression

Visual opsins represent only a small proportion of the genes expressed in the retina. To gain a broader understanding of light processing within the retina, we explored wider patterns of differential expression; comparative transcriptomic analysis revealed considerably fewer genes differentially expressed overall between the grouper geminates (6.6%) as compared with the damselfish geminates (17.7%). Such differences may reflect differences in the timing of differentiation between the damselfish and grouper geminates, with recent estimates based on genome-wide patterns of divergence indicating that the damselfish geminates likely diverged considerably earlier (∼12 million years ago) than the grouper geminates (∼5 million years ago) (Tysall et al. 2026).

We focused more in-depth analyses of expression patterns on three key sets of genes likely to be most impacted by differences in the underwater light field, specifically, those involved in (a) phototransduction cascade, (b) circadian clock regulation, and (c) non-visual opsins.

#### ***a.*** Phototransduction cascade

The broader spectral bandwidth and higher underwater light levels in the CAR compared to TEP suggest that genes in the cone phototransduction pathway should be overexpressed in the CAR geminates, whilst the rod phototransduction pathway should be more active in the TEP geminates (in line with the overexpression of *RH1* dim-light opsins discussed above).

In line with the first expectation, we find that genes with significantly different expression in the cone pathway tended to be overexpressed in the CAR geminate (Fig. 3; Tables S7-S8). This effect was particularly noticeable in the damselfish, where all five significant cone specific phototransduction genes (including *GNGT2*, *PDE6C, PDE6H, GRK7* and *ARR3*) exhibited this pattern (Fig. 3A; Table S7). Comparatively, in the groupers, only two cone genes showed significant differences, *GNAT2* and *GUCA1C,* but both were in the expected direction (Fig. 3B; Table S8). As was the case for the opsins, different genes were involved in the adaptation of the two groups to different light conditions.

**Figure 3.**
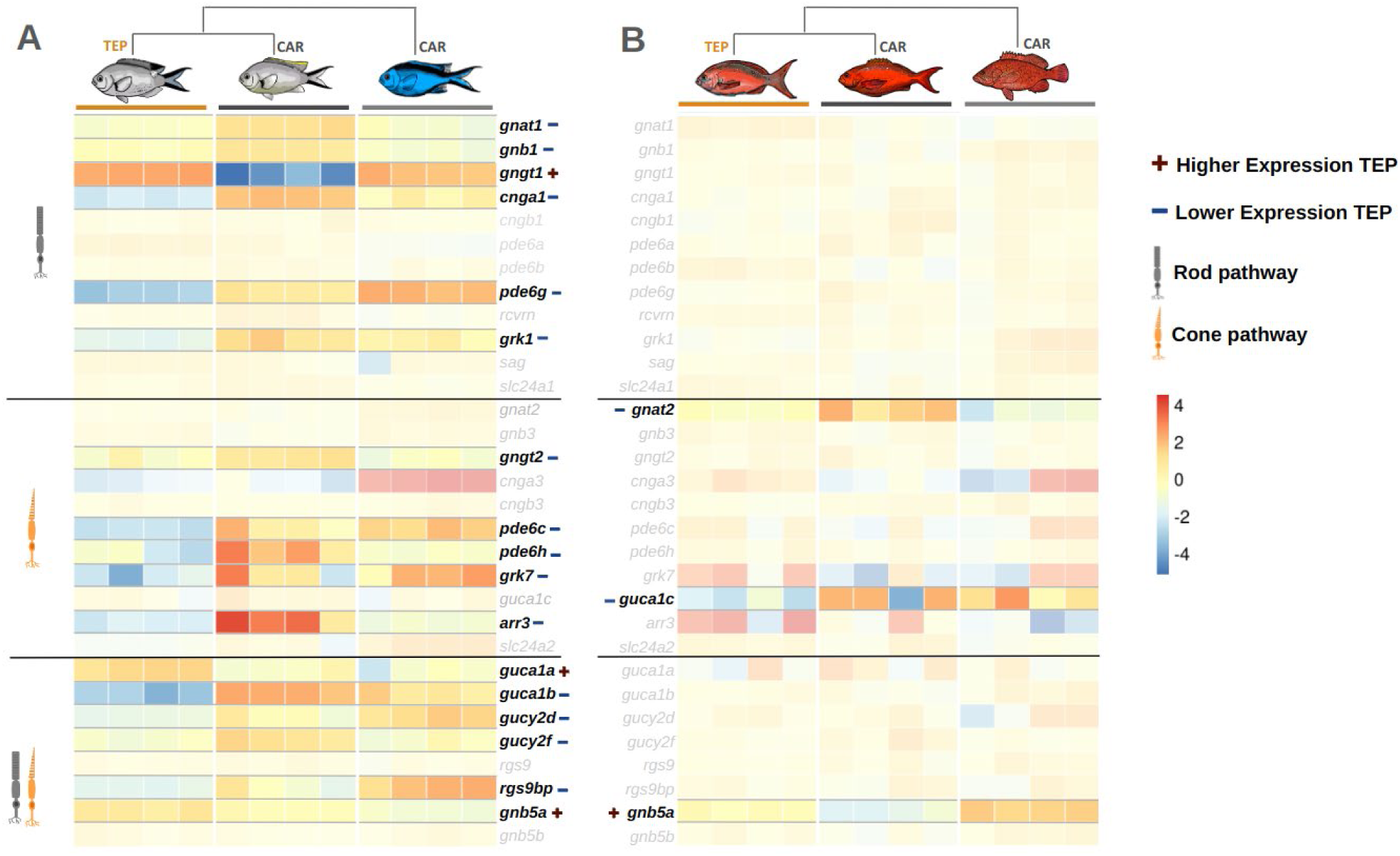
Heatmap of variance-stabilising transformed read counts for phototransduction cascade related genes in retinas from four individuals per species: **A)** the damselfish and **B)** the groupers geminate pairs and their respective outgroup species. Those significantly differentially expressed between the TEP and Caribbean geminate species are highlighted. The (+) and (-) symbols indicate whether the expression in the TEP geminate is significantly higher or lower than that of the Caribbean geminate. Rod specific pathway genes, cone specific pathway genes and those utilised by both pathways are indicated.

**Figure 4.**
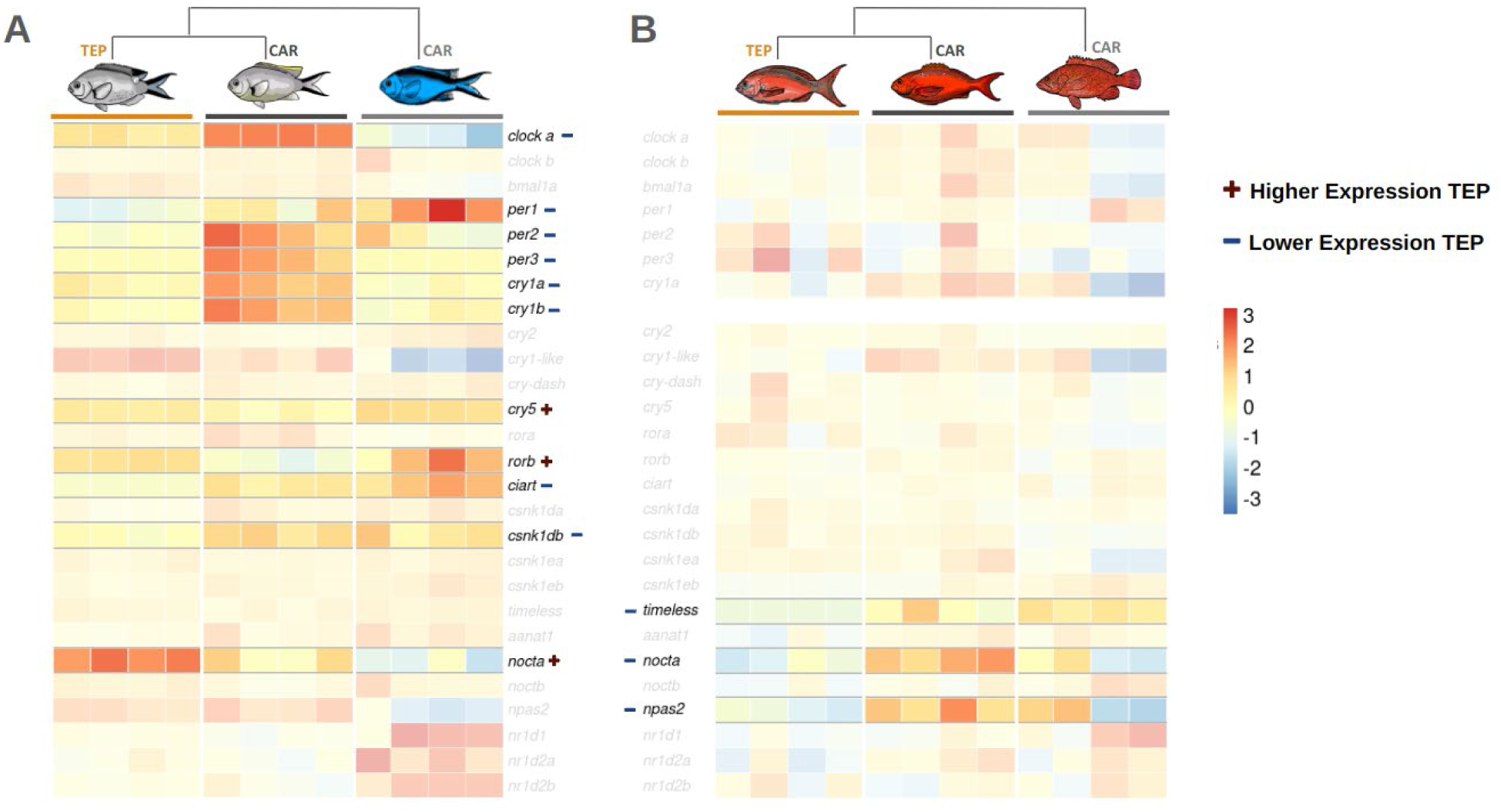
Heatmap of variance-stabilising transformed read counts for circadian related genes in retinas from four individuals per species: **A)** the damselfish and **B)** the groupers geminate pairs and their respective outgroup species. Those significantly differentially expressed between the TEP and Caribbean geminate species are highlighted. The (+) and (-) symbols indicate whether the expression in the TEP geminate is significantly higher or lower than that of the Caribbean geminate.

**Figure 5.**
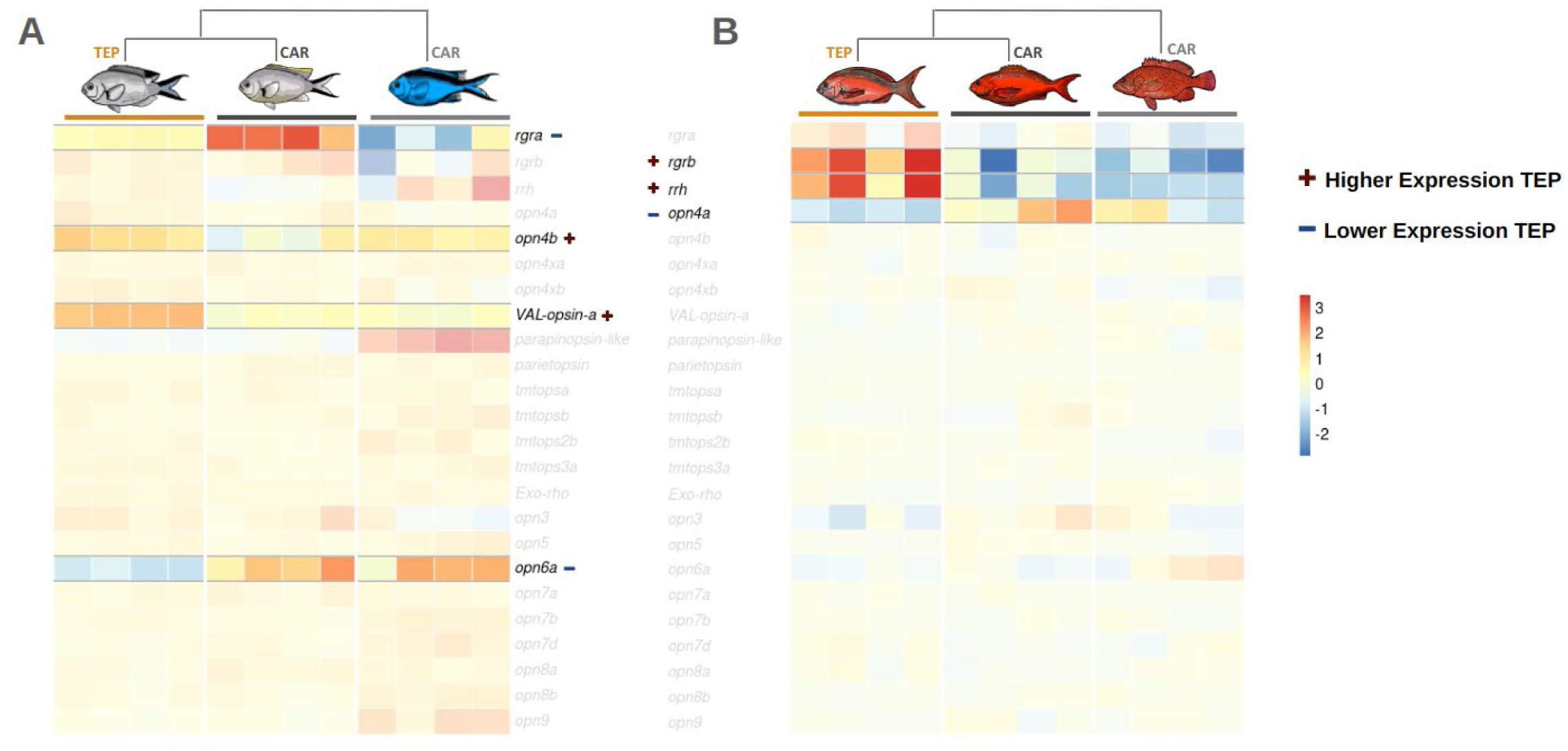
Heatmap of variance-stabilising transformed read counts for non-visual opsins in retinas from four individuals per species: **A)** the damselfish and **B)** the groupers geminate pairs and their respective outgroup species. Those significantly differentially expressed between the TEP and Caribbean geminate species are highlighted. The (+) and (-) symbols indicate whether the expression in the TEP geminate is significantly higher or lower than that of the Caribbean geminate.

For the rod pathway, the results were less clear cut. Whilst we did not find any differences for the groupers, most of the rod pathway genes detected in the damselfish were overexpressed in the CAR geminate; thus, matching the pattern found in the cone pathway, but not matching our *a priori* expectation. Indeed, only one rod pathway gene (*GNGT1)* showed overexpression in the TEP (Fig. 3A). A further six non-rod or cone specific phototransduction genes were differentially expressed between the damselfish geminates, with the majority also more highly expressed in the CAR geminate (*GUCA1B*, *GUCY2D*, *GUCY2F* and *RGS9BP*). However, two, *GUCA1A* and *GNB5A,* showed the opposite pattern, with the latter differentially expressed in the same direction in the groupers.

#### ***b.*** Circadian regulation genes

We anticipated that the differences in levels of underwater light available and spectral content between the two basins might affect the entrainment of circadian rhythms, both directly in response to the characteristics of the spectral field and indirectly through their influence on the effective daylength at depth (Collin and Hart 2015).

Meeting this expectation, in the damselfish, we found a predominance of overexpression in the CAR geminate, spanning core clock component genes central to the transcription-translation feedback loop; *CLOCKA*, *period* genes (*PER1*, *PER2* and *PER3*), *cryptochrome* genes (*CRY1A* and *CRY1B*), as well as accessory loop clock genes *CIART* and *CSNK1DB* (Fig. 4A; Table S7). Circadian related genes differentially expressed in the opposite direction (more highly expressed in the TEP geminate) include downstream effector *NOCTA,* accessory loop regulator *RORB* and further light induced cryptochrome *CRY5*.

In the groupers, we found a similar pattern, overexpression of core circadian genes in the CAR geminate; however, the genes involved were fewer and, largely, different. All the differentially expressed circadian related genes we recovered were more highly expressed in the CAR geminate, namely, *TIMELESS* (a direct modulator of the *Drosophila* circadian clock, though its role in teleosts is not well characterised (Cai and Chiu 2022)), *NPAS2* and *NOCTA* (Fig. 4B; Table S8). *NPAS2* (like *CLOCKA* which is differentially expressed in the damselfish) is part of the *CLOCK* gene family and has a role in the core circadian clock as part of the positive regulatory arm of the main feedback loop. Finally, *NOCTA,* the only circadian gene differentially expressed in both the damselfish and in the grouper geminate pairs, is interestingly differentially expressed in opposite directions.

#### ***c.*** Non-visual opsins

Non-visual opsins are photo-sensitive genes with a diverse range of non-image forming functions in the retina. Since we expect different light environments to result in different physiological and behavioural responses, we predicted non-visual opsins utilised in such responses may be differentially expressed in the contrasting light environments of the TEP and Caribbean (but the direction would depend on the exact function of each gene).

We found several non-visual opsins involved in the retinoid cycle, which regenerates the visual pigment used by photoreceptor cells to be differentially expressed between the geminates of both families (Fig. 5; Tables S7-S8). We expected that the higher light levels characterising the underwater habitat of our CAR geminate members would cause overexpression of those genes involved in regeneration of the visual pigment compared with their TEP counterparts. This expectation held true for the retinal G protein coupled receptor teleost paralog *RGRA*, which was more highly expressed in the damselfish CAR geminate, however in the groupers the opposite pattern was present with paralog *RGRB* and the retinal pigment epithelium-derived rhodopsin homolog (*RRH*), thought to be involved in the storage of *all trans* retinol within the RPE, actually exhibiting higher expression in the grouper TEP geminate member (Fig. 5).

Melanopsins are thought to play a role in both entrainment of circadian rhythms and other non-visual tasks such as pupil-response in mammals (Panda et al. 2002, Lucas et al. 2003). In zebrafish, all five *OPN4* variants have been found to be expressed in the retina with non-overlapping patterns of expression and apparent functional diversity (Davies et al. 2011). We found different melanopsin variants differentially expressed between the damselfish and grouper geminates in different directions potentially reflecting this functional diversity. Melanopsin variant *OPN4B* showed increased expression in the damselfish TEP geminate, whereas in the groupers melanopsin variant *OPN4A* showed the opposite pattern of expression being less expressed in the TEP geminate (Fig. 5; Tables S7-S8). Two additional non-visual opsins were differentially expressed between the damselfish geminates and not found in the groupers. Neuropsin *OPN6A,* known to be expressed in photoreceptors in zebrafish (Davies et al. 2015, Man et al. 2023), was downregulated in the TEP damselfish geminate (Fig. 5A; Table S8). The vertebrate ancient long opsin (*VAL-opsin a*), thought to be involved in light-dependent control of gap junctions or phase setting of the retinal circadian clock in zebrafish (Kojima et al. 2000), was upregulated in the TEP geminate of the damselfish (Fig. 5A; Table S7).

## Discussion

We examined the divergence in the light environment and visual system of young pairs of allopatric sister taxa (trans-isthmian geminate species) living in either a stable, oligotrophic and blue-shifted sea, the Caribbean (CAR), or in the nutrient-rich, turbid and long wavelength-shifted Gulf of Panama, Tropical East Pacific (TEP). We found that, despite their phylogenetic divergence, each species pair underwent similar phenotypic adaptations in response to TEP or CAR photic environments. The compression of the underwater light field from CAR to TEP waters, particularly pronounced at the shortest wavelengths, was tracked by parallel shifts in visual sensitivity towards the centre of the underwater light spectrum in both TEP representatives of *Azurina* damselfish and *Cephalopholis* groupers.

By teasing apart the contributions to the predicted maximal sensitivity of single (shorter wavelength) cones and double (longer wavelength) cone expression (Carleton et al. 2008), we uncovered a pattern of phenotypic parallelism in the relative response of single and double cones to a shift in photic environment. Remarkably, such parallel, coordinated adaptation of cone type expression involved phylogenetically distant lineages of teleosts.

In both TEP damselfish and TEP grouper species, short-wavelength cone sensitivities are shifted to longer wavelengths, while long-wavelength cone sensitivities move to shorter wavelengths compared to their Caribbean counterparts; however, such changes are accomplished by different opsins in the two geminate pairs. Non-parallelism at the molecular level is perhaps to be expected between phylogenetically distant lineages, carrying different sets of opsin classes and their paralogs. In addition, different sets of opsins in different lineages can potentially accomplish similar visual functions. This is because the same spectral region can be occupied by opsins belonging to different classes, i.e. the ranges of maximum sensitivity of different opsin classes partially overlap (Bowmaker 2008). Finally, the potential for co-expression of multiple opsins within a single photoreceptor cell, as documented in some mammals and fishes (Lukáts et al. 2002, Jacobs et al. 2004, Dalton et al. 2014), as well as in this study, creates de facto new photoreceptor classes sensitive to intermediate wavelengths, further expanding the range of possible solutions in response to similar photic environments.

In the damselfish geminate pair, the effect of the different underwater light environments extended beyond visual opsins, to affect a large set of genes in the retina, particularly those in the downstream phototransduction cascade. It is through this signalling cascade that light absorbed by a photoreceptor cell is converted into an electrical signal (Pugh et al. 1999, Pugh and Lamb 2000, Lamb et al. 2016, Lamb and Hunt 2017). The expression of genes in the phototransduction cascade are likely therefore to be pertinent to the rate of activation, deactivation and recovery of opsins and in turn the speed and sensitivity of photoresponse (Pugh et al. 1999, Arshavsky et al. 2002, Umino et al. 2012). The striking pattern of greater expression of a large swathe of phototransduction genes in the Caribbean damselfish, prominent in both rod and cone pathways, may reflect the need to prioritise the speed of photoresponse in the brighter light environment of the Caribbean. In contrast, lower expression of phototransduction genes, possibly leading to slower refresh rate of visual information in the TEP geminate, may allow for greater visual sensitivity in the dimmer light conditions typical of the TEP. While only a limited number of studies of visual adaptation to underwater light currently extend beyond opsins to changes in the downstream phototransduction pathway, there is evidence for environmentally relevant shifts in this pathway in other systems. Multiple phototransduction pathway genes have been found to be downregulated in the Japanese medaka, under shorter-day (reduced light) seasonal conditions (Shimmura et al. 2017). Evidence for selection has been found in the coding sequence of several phototransduction genes, in response to depth adaptation in Malawi cichlids (Malinsky et al. 2018). We focus here only on one mechanism of adaptation, namely, gene expression, but changes in coding sequence may also be important, as shown in cichlid fish (Malinsky et al. 2018). The much-reduced pattern of differential expression occurring in the groupers may reflect different genetic mechanisms, as just mentioned, and/or underlie finer-scale differences in adaptive responses of the visual system to the underwater light field.

Differences in the spectral content and levels of underwater light available to CAR or to TEP members of each geminate pair appear to have affected the entrainment of circadian rhythms, either directly in response to the characteristics of the spectral field, or indirectly, through the influence of turbidity on the effective daylength at depth (Collin and Hart 2015). We again found these effects most prominently in the damselfish, where almost all elements of the primary transcription-translation feedback loop of the circadian clock (*CLOCKA*, *PER1*, *PER2*, *PER3*, *CRY1A* and *CRY1B*) were more highly expressed in the Caribbean geminate. Notably, *CRY1A* and *PER2* have been identified as playing important roles in conveying light information to the core clock in zebrafish, their expression increasing in response to light and, in the case of *CRY1A*, inducing phase shifts (Ziv et al. 2005, Tamai et al. 2007, Frøland Steindal and Whitmore 2019). Groupers followed a similar pattern, though different genes were often involved. For example, *NPAS2*, a paralog of *CLOCKA*, known to modulate contrast sensitivity, at least in mammals (Reick et al. 2001, DeBruyne et al. 2006, 2007, Hwang et al. 2013), was also more highly expressed in the Caribbean geminate. On the other hand, *NOCTA*, a gene involved in metabolism regulation and with a rhythmic expression peaking at night (Green and Besharse 1996, Green et al. 2007, Estrella et al. 2019), was differentially expressed in both families, but with opposite differential expression patterns. Circadian-related genes featured prominently in a study of differential gene expression between deep and shallow water cichlids (Musilova et al. 2019), including some identified here (*PER1*, *RORB*). However, research on circadian regulation and entrainment in response to underwater light of different spectral quality or intensity is still in its infancy (Piechura et al. 2017). So far, studies have been limited to very few species and it is unclear whether individual genes and their paralogs play the same coordinated role across taxa or whether conservation of feedbacks and other regulatory circuits is not strictly replicated at the individual gene level. These gaps in our understanding of circadian regulation make gene-by-gene comparisons of the effects of a particular spectral stimulus, between distantly related lineages, as is our case, at best challenging, if not problematic, to interpret.

Non-image forming sensitivity to light is often mediated by non-visual opsins with effects on a broad range of behavioural and physiological responses (Davies et al. 2015). The melanopsins and other non-visual opsins involved in the retinoid cycle showed differential expression between Caribbean and TEP geminates of both families. This suggests coordinated tuning of light-sensitive factors to directly affect vision (e.g. *RGR*). The opposing patterns of expression in two different melanopsins in the damselfish (*OPN4B*) and groupers (*OPN4A*) may be explained by the non-overlapping patterns of expression, differences in their distribution, and apparent functional diversity of melanopsins found in the zebrafish (Davies et al. 2011). This diversity in function and expression has led to their proposed role in allowing finer scale adjustments in the retina, to respond to highly varied light environments such as the marine habitats investigated here (Davies et al. 2011).

### UV sensitivity (or the lack thereof) in the planktivore groupers

The lack of expression of the UV-sensitive *SWS1* opsin present in the planktivorous groupers geminates was surprising, considering the proposed role of UV vision in plankton detection (Browman et al. 1994, McFarland and Loew 1994, Jordan et al. 2004, Rick et al. 2012, Novales Flamarique 2013) and given the expression of *SWS1* in the geminate damselfish, foraging on the same reefs on a similar set of plankton species, including those with the smallest minimum sizes (Randall 1967; Fig. S1). The presence of UV-sensitive *SWS1* opsins, even in non-planktivorous fishes, is rather common, given that early life stages typically forage on plankton; in species where adults transition to other foraging niches, *SWS1* is later replaced by *SWS2* expression, as observed for example in another serranid, the longtooth grouper (*Epinephelus bruneus*) (Matsumoto and Ishibashi 2015), or in salmonids whose single cones irreversibly switch expression from *SWS1* to *SWS2* in the juvenile phase, when plankton feeding is abandoned (Cheng and Novales Flamarique 2004, Cheng et al. 2006, Cheng and Flamarique 2007). Given the recent emergence of adult plankton-feeding in the *Cephalopholis* geminates lineage and the advantage provided by UV vision in other planktivorous species, we would have expected *SWS1* expression in these species: however, the eye lens in *Cephalopholis furcifer* is not UV-transparent in the adult (Fig. S15), in line with the lack of *SWS1* expression in its retina, and suggesting that UV vision is not a prerequisite for foraging on plankton of the same sizes and species targeted by the UV-sensitive damselfish in this study (Randall 1967; Fig. S1). It is possible that the evolution of planktivory in the *C. furcifer/colonus* pair might be recent and this lineage might be retaining an ancestral UV-absorbing adult lens, typical of many groupers, including the geminates’ closest extant relative, *C. fulva* (Pierotti, *unpubl. data*).

While *C. furcifer* (and likely *C. colonus*) are clearly capable of obtaining the very same species and size classes of plankton that the UV-sensitive *Azurina* usually catch, UV sensitivity might provide the latter with superior detection efficiency and/or increased sighting distance.

## Conclusions

Our study highlighted parallel adaptation of the visual system of two distant groups of coral reef fishes to either bright, broad-spectrum conditions as found over Caribbean reefs or comparatively dimmer, red-shifted and narrower spectrum of underwater light in the Gulf of Panama (Tropical East Pacific). Both the *Azurina* damselfish and the *Cephalopholis* grouper ‘geminates’ tune their visual system to maximise light catch and their visual sensitivity but this is achieved by recruiting and modulating the expression of different sets of opsin genes, as expected given the lineage-characteristic opsin toolsets. In addition to differential opsin expression, amino acid replacements at known tuning sites and co-expression of different opsins within the same photoreceptor cell led to visual systems optimally tuned to the local prevailing light spectrum. Patterns of differential expression in other genes involved in light response suggest the impact of diverging underwater conditions between Caribbean and TEP might extend well beyond vision. Indeed, the closure of the Isthmus of Panama brought about differences in salinity, nutrients and temperature (Haug and Tiedemann 1998, D’Croz and O’Dea 2007) that likely imparted strong selective pressures on marine organisms on either side of the Isthmus of Panama. With the emergence of multiple pairs of geminate species, this represents an ideal natural experiment at global scales and an exceptional opportunity for future studies of rapid, parallel evolution at the level of single phenotypic traits, whole genomes and entire communities.

## Materials and Methods

### Underwater spectral measurements

In order to characterise the underwater photic environment of collecting sites (Piscadera Baii, Curacao Island, Southern Caribbean Sea; Isla Saboga, Gulf of Panama, Tropical Eastern Pacific), we measured spectral irradiances from downwelling, upwelling and sidewelling underwater light, just under the surface and at 2.5m, 5.0m and 7.5m depth, as described in Pierotti et al. (2020) (Fig.S2). We took all measurements between 11.00 and 13.00 on a sunny clear day.

We described the spectral distribution of underwater irradiance using two metrics, i. the wavelength λP50 at which the total number of photons in the underwater light spectrum is equally divided (Munz and McFarland 1973), and ii. the width of the available light spectrum expressed as the difference Δλ between λP25 and λP75 cumulative photon frequencies, across the range 350-650nm. We also calculated the total number of photons available at each depth at the two sites as photon flux density at depth (Table S1).

### Satellite data

Level-3 (4 km resolution) monthly mean remote sensing reflectance (R_rs_(λ)) data from the MODIS-Aqua sensor were obtained from NASA’s Ocean Biology Distributed Active Archive Center (OB.DAAC; https://oceancolor.gsfc.nasa.gov/) at six wavelengths (λ = 412, 443, 488, 531, 547, and 667 nm) for the calendar years 2010 – 2019. Twelve coral reef sites in the Tropical East Pacific and the Caribbean Sea were chosen to represent regional variation in these two basins (Fig. S3). Monthly mean R_rs_(λ) values were extracted from each site, and every year, in the months of February (the dry season, typically a period of high primary productivity associated with upwelling events in the Gulf of Panama) and September (the wet season, characterised by lower levels of primary productivity), using bilinear interpolation.

### Remote sensing-based estimation of downwelling irradiance at depth

Inherent optical properties (IOPs) describing the total absorption and backscattering characteristics of seawater at reference sites were then derived from the MODIS R_rs_(λ) inputs, following the Quasi-Analytical Algorithm (QAA) of Lee et al. (2002), and then expanded from discrete MODIS bands to hyperspectral resolution (spectrally continuous bands at 10 nm intervals from 400–700 nm), following the methods outlined in Lee et al. (2021) and Lee et al. (2005b). The hyperspectral IOPs were subsequently used as model inputs to analytically derive estimates of spectrally resolved diffuse attenuation coefficients (K_d_(λ), Lee et al. 2005a, 2021). The K_d_(λ) terms describe the exponential slope of light extinction as a function of depth, enabling the estimation of spectral diffuse irradiance at any depth, E_d_(λ,z), following the Beer-Lambert Law:

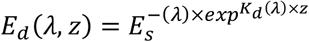

where E_s_^-^(λ,z) is the spectral downwelling irradiance just below the sea surface. Plainly, these coefficients enable the reconstruction of the sub-surface light field based on the optical properties estimated from satellites. Finally, for each site and date, E_d_(λ,z) spectra were propagated to depths of 1, 8, 16, and 24 feet (0.3048, 2.438, 4.876, and 7.314 meters), conditioned on the site-specific solar zenith angle occurring at local noon (12:00 local standard time) on February 15 and September 15 (computed using the R package solrad).

Additionally, cross-sectional visualisations of the underwater light field were generated across the twelve sites in the TEP and CAR. Spectrally resolved downwelling irradiance profiles were propagated continuously from the surface to 100 m depth at 2 m intervals using the satellite-derived Kd (λ) fields and the Beer–Lambert relationship. At each wavelength, irradiance values were normalised to the corresponding modelled surface irradiance (Ed(λ,z)/Ed(λ,0)), such that pixel brightness represented the fraction of surface light remaining with depth, while hue was assigned according to wavelength-based RGB colour mapping. This approach enabled qualitative visualisation of both the spectral composition and relative intensity of the underwater light field across optically distinct water masses.

### Remote sensing-based spectral metrics

To facilitate comparisons between sites and seasons, details of the water colour were summarized using the Apparent Visible Wavelength (AVW) metric (Vandermeulen et al., 2020). The AVW condenses the full spectral distribution of R_rs_(λ) (or modelled irradiance) into a single wavelength value, providing a compact measure of spectral shifts related to water optical properties. Generally, lower AVW values (440 - 490 nm) represent relatively clear, less productive waters while higher AVW values (490 - 600 nm) represent more turbid, productive water masses. We analysed apparent visible wavelength (AVW) at 30 m depth using a linear mixed effects model (LMM) fitted with the lmer function in the R package lme4 (Bates et al. 2015). Region (Caribbean, CAR; Tropical Eastern Pacific, TEP) and season (dry, wet) were included as fixed effects together with their interaction, and site as a random intercept. An initial model including year as an additional random intercept yielded zero variance for that term and was therefore dropped. Fixed effect significance was assessed using Type III *F*-tests with Kenward-Roger approximation for denominator degrees of freedom, as implemented in the lmerTest package (Kuznetsova et al. 2017). Post-hoc pairwise contrasts between regions within each season, and between seasons within each region, were computed using the emmeans package (Lenth 2024). Bootstrap confidence intervals (parametric) for model coefficients were additionally computed using the confint function to provide robust interval estimates for random effect variance components and for fixed effect terms not directly available from the pairwise contrasts (Davison & Hinkley 1997). Model assumptions were verified by visual inspection of residual plots (residuals vs. fitted values and Q-Q plots of standardised residuals), both overall and by site. All analyses were conducted in R v4.x (R Core Team 2024).

### Study site and sampling

We collected fish with SCUBA at depths of between 3-20m over or just above coral reefs, at various sites along the Western (leeward) coast of the island of Curacao, Southern Caribbean, and on reefs around Isla Saboga, in the Islas Perlas archipelago, Tropical Eastern Pacific. Individual fish were chased into 150cm x 60cm barrier nets, then kept in large, black baskets secured with a scuba mesh bag during underwater collection and transfer to the surface. We euthanized fish immediately with an overdose of buffered MS-222 followed by decapitation. We placed muscle tissue for DNA extraction in 96% ethanol and moved it to a - 80°C freezer upon arrival at the research facilities, always within three hours from collection. Additionally, eyes were enucleated and the retina was extracted and placed in RNAlater solution within 10 minutes of euthanasia.

### Micro-spectrophotometry of photoreceptor cells

We performed retina micro-spectrophotometry (MSP) on adult individual fish within 24hrs from collection in the field. Fish were maintained under dark conditions for 2 hours prior to MSP and then euthanized with an overdose of MS-222, followed by cervical dislocation. The eyes were rapidly enucleated under dim red light, and the retinas extracted while in a buffer solution (PBS with 6% sucrose, pH 7.2). Small pieces of the retina were separated from the retinal pigment epithelium, placed on a cover slide and fragmented, using razor blades and tungsten needles, to isolate individual photoreceptors. The preparation was sealed with a second cover slide and Corning High Vacuum grease.

We used a single-beam, computer-controlled MSP, with a 100 W quartz iodine lamp that allowed for accurate absorption measurements down to 340 nm (Loew 1994). The wavelength of maximum absorbance (*λmax*) of an individual photoreceptor cell was obtained by fitting templates to the interpolated, normalised absorbance spectra (Lipetz and Cronin 1988, Govardovskii et al. 2000) with criteria for data inclusion following Loew (1994). MSP records were processed as in Escobar-Camacho et al. (2019) and Pierotti et al. (2020). Following Dalton et al. (2014), absorbance curves from cones coexpressing two opsins were analysed with visual pigment templates for mixtures of two pigments, by setting the *λmax* of the contributing pure pigments (as measured in “pure” cones) and best fitting the measured absorbance with least squares by varying the relative proportions of each pigment in the mixture.

### RNA extraction and sequencing

We extracted total RNA from *n* = 4 individuals per species with a RNeasy kit (Qiagen), quantified and quality checked on an Agilent 2100 Bioanalyzer. We prepared RNAseq libraries using a TruSeq RNA Sample kit (Illumina, San Diego) at the Naos Marine Labs, Smithsonian Tropical Research Institute, Panama. We barcoded individual species and multiplexed twelve to a lane of an Illumina HiSeq 2000 to generate 150bp paired-end sequences.

### Quality control and de novo assembly

We initially evaluated the quality of the raw RNA-seq reads using FastQC v0.11.9 (http://www.bioinformatics.bbsrc.ac.uk/projects/fastqc/). Following this, we filtered and removed any residual ribosomal RNA (rRNA) contamination in the reads using SortMeRNA v4.2.0 (Kopylova et al. 2012). We used Cutadapt v3.2 (Martin 2011) to remove adaptor sequences, quality trim reads and remove reads under a minimum read length threshold of 30kb. We quality trimmed reads by removing those with a phred score below 20. However, we used a less stringent quality cutoff of <5 in the transcriptome assembly pipeline as excessive quality trimming can negatively impact *de novo* assemblies (MacManes 2014). As part of the *de novo* assembly pipeline, we also used rCorrector v1.0.4 (Song and Florea 2015), to correct any very rare, and therefore likely erroneous, K-mers to more frequently occurring K-mers, in order to reduce the impact of erroneous K-mers on transcriptome assembly. After each filtering step, we re-assessed read quality using FastQC and evaluated the overall change in quality, following the quality control pipeline using MultiQC v1.9 (Ewels et al. 2016).

We combined reads from all four replicate individuals to construct a *de novo* transcriptome assembly for each species. To assemble the transcriptomes, we used Trinity v2.13.2 (Haas et al. 2013) on default settings. Following transcriptome assembly, we used BUSCO (Benchmarking Universal Single-Copy Orthologs) v5.1.2 (Simão et al. 2015) to assess transcriptome quality, where the proportion of complete BUSCOs is indicative of transcriptome completeness, based on the Actinopterygii odb9 database. We carried out further assessment of transcriptome quality through the comparison of basic transcriptome contig metrics, including contig N50, i.e. the smallest contig length that includes 50% of assembled transcript nucleotides. In addition, we computed the contig ExN50 statistic to evaluate the contig structure of our assemblies. In comparison to the traditional contig N50 statistic, the ExN50 statistic is more appropriate for transcriptome assemblies as it takes into account transcript expression. We therefore computed the N50 statistics but limited it to the most highly expressed transcripts representing x% of the expression data, using Trinity toolkit utilities scripts ‘align_and_estimate_abundance.pl’, ‘abundance_estimates_to_matrix.pl’ and ‘contig_ExN50_statistic.pl’.

### Functional annotation and transcriptome expression quantification

Transcriptomes were functionally annotated with eggNOGG-mapper v2.1.5 (Huerta-Cepas et al. 2019, Cantalapiedra et al. 2021) with default settings, and the flag --tax_scope 7898 to restrict the taxonomic scope used for annotation to the Actinopterygii. In order to contrast differential expression of the same genes between sister species, we only used gene annotations which were present in all three species (geminate pair + outgroup), as a comparison set in the differential expression analysis. We quantified transcript expression using pseudo-aligner Salmon v1.9.0 (Patro et al. 2017) with the additional flags: --seqBias, --gcBias and –posBias, to enable Salmon to correct for any random hexamer priming bias, fragment-level GC biases and non-uniform coverage biases in the RNA-seq data (Patro et al., 2017). A high proportion of reads (between 95.9% - 97.7%) mapped back to their respective transcriptome assembly following Salmon transcript expression quantification.

In the damselfish set, we recovered 14,346 genes that were shared between all three species and used in the differential expression analysis. In the grouper set, we recovered and took forward 15,009 shared annotated genes.

### Gene extraction and phylogenetic reconstruction

Using the MAFFT v1.5.0 (Katoh et al. 2002) alignment plugin within Geneious Prime 2026.0.2 (https://www.geneious.com), we confirmed the identity and class of putative opsin gene sequences, and other differentially expressed genes of interest, by visualising and aligning the sequences back to multiple teleost sequences obtained from GenBank (www.ncbi.nlm.nih.gov/genbank/). To assess the lineage and opsin class of each visual opsin sequence, we used the nucleotide coding region of sequences from both our damselfish and grouper study species, alongside a phylogenetically diverse range of teleost opsin sequences obtained from GenBank and constructed maximum likelihood gene trees using the RAxML v4 (Stamatakis 2006) plugin in Geneious Prime. We used the nucleotide model GTR Gamma, ‘Rapid bootstrapping and search for best-scoring ML tree’ algorithm and undertook 50 bootstrap replicates. Accession numbers for all sequences used in each opsin gene tree are provided in supplementary Table S2.

### Coding sequence variation

To uncover potential shifts in opsin spectral sensitivity, we searched for amino acid substitutions in the translated opsin coding sequences that fell in sequence regions corresponding to putative transmembrane regions, the retinal binding pocket, or known spectral tuning sites. Damselfish and grouper sequences of each opsin class were aligned to bovine rhodopsin *RH1* (ACCESSION: NM_001014890) to identify tuning site position. The location of retinal binding pockets and transmembrane regions were taken from Hofmann et al. (2012).

### Maximal spectral sensitivity

To simultaneously evaluate the effects of both cone spectral shifts and changes in opsin expression in each cone type on overall retinal sensitivity across lineages, we calculated the predicted maximal spectral sensitivity index PS_max_ (Carleton et al. 2008, Hofmann et al. 2009). Average single-and double-cone sensitivities were obtained by weighing peak spectral sensitivities for each opsin by the fraction of their expression in each cone type. In the case of cones co-expressing two opsins, we assigned relative contributions as follows. In the absence of information on the relative frequency of such cones in the retina, we considered equal frequencies of each cone type (pure and coexpressing); we then assumed the relative composition in co-expressing cone types to be as predicted from micro-spectrophotometry.

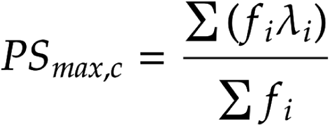

where C is either the single-or the double-cone photoreceptor type, f_i_ is the percent expression of the *i*th opsin out of the total, and λ_i_ is the corresponding peak absorbance of that opsin in the species.

### Relative cone opsin gene expression analysis

Transcripts Per Million (TPM) counts were extracted from the Salmon quantification output. To obtain the proportion of each cone opsin class expressed by an individual, each cone opsin was normalised by the total cone opsins counts. We used the same procedure to calculate relative expression for opsins specifically expressed in either double or single cones.

### Transcriptome wide differential expression analysis

We conducted the differential expression analysis in R v4.2.1 (R Core Team, 2021), using the DESeq2 package v1.36.0 (Love et al. 2014). We summed isoform level count estimates to gene level using the R package Tximport (Soneson et al. 2016) which corrects for differences in average transcript length across samples. We identified a predefined set of genes in the retina relating to the phototransduction cascade, circadian clock regulation and non-visual opsins whose expression may be directly affected by differences in light environment, and for which the automated gene annotation could be manually checked against other related species again by visualising and aligning the sequences back to multiple teleost sequences obtained from GenBank (www.ncbi.nlm.nih.gov/genbank/). Adjusted *p*-values were calculated based only on this subset. We required differentially expressed genes to show a minimum log2 fold change of 0.5 (*lfcThreshold = 0.5*) and Benjamini-Hochberg (BH)-adjusted *p*-value <0.05 (*alpha = 0.05*) between the geminate species. While we were primarily interested in genes differentially expressed between the CAR and TEP members of each geminate pair, we additionally carried out pairwise differential expression analysis between the Caribbean geminate and the (CAR) outgroup in each family, to help contextualise findings.

