## Supplementary Materials for "Parallel phenotypes underpinned by different genes in the visual system of two trans-isthmian coral reef fish species pairs"

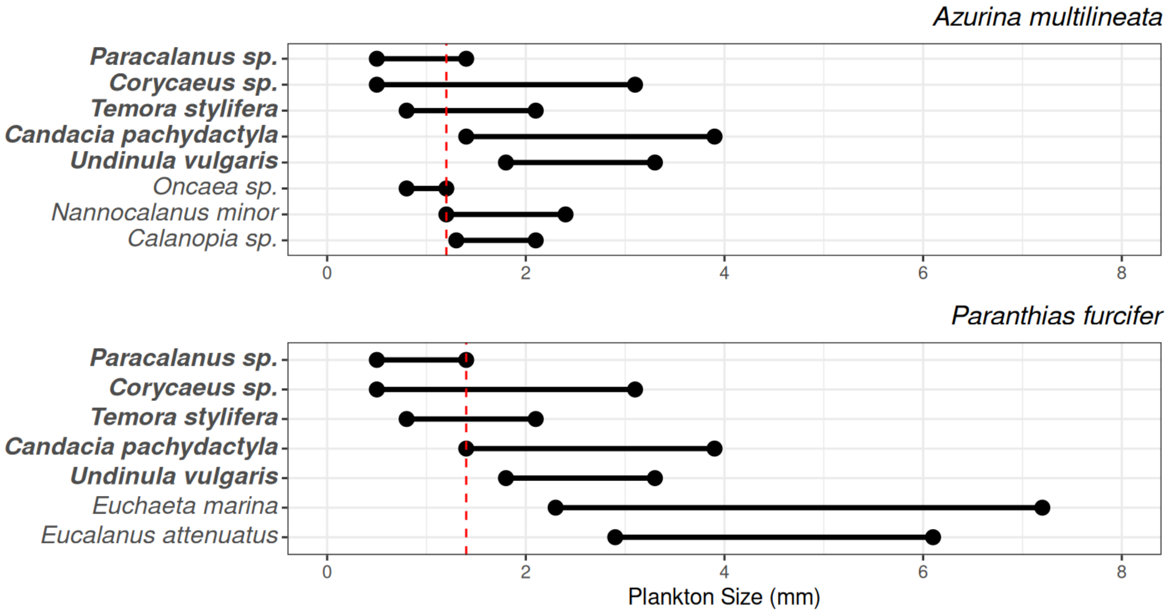


**Figure S1.** Plankton species, and their size range, commonly found in the diet of the damselfish *A. multilineata* and the serranid *Cephalopholis furcifer*, according to Randall (1967). *In bold*, species shared by both damselfish and grouper. If a plankton species is recorded in the stomach content of a fish species, then the size of that plankton item is most likely below its maximum recorded from the literature. That allows to identify a minimum prey size (i.e. the largest size class recorded) that the fish species can eat (*in red*).

**Figure S2**. **A**. In situ measurements of downwelling (*left*), average sidewelling (*centre*), and upwelling (*right*) irradiances, on a vertical depth profile, just under the surface (—), at 2.5m (—), 5.0m (---) and 7.5m (···). **B**. depth-dependence of the prevalent wavelength $\lambda P_{50}$, and **C**. depth-dependence of the spectral bandwidth $\Delta\lambda P_{50}$, of the downwelling, sidewelling and upwelling irradiances (Curacao, Caribbean in *blue*, Isla Saboga, Tropical East Pacific in *green*).

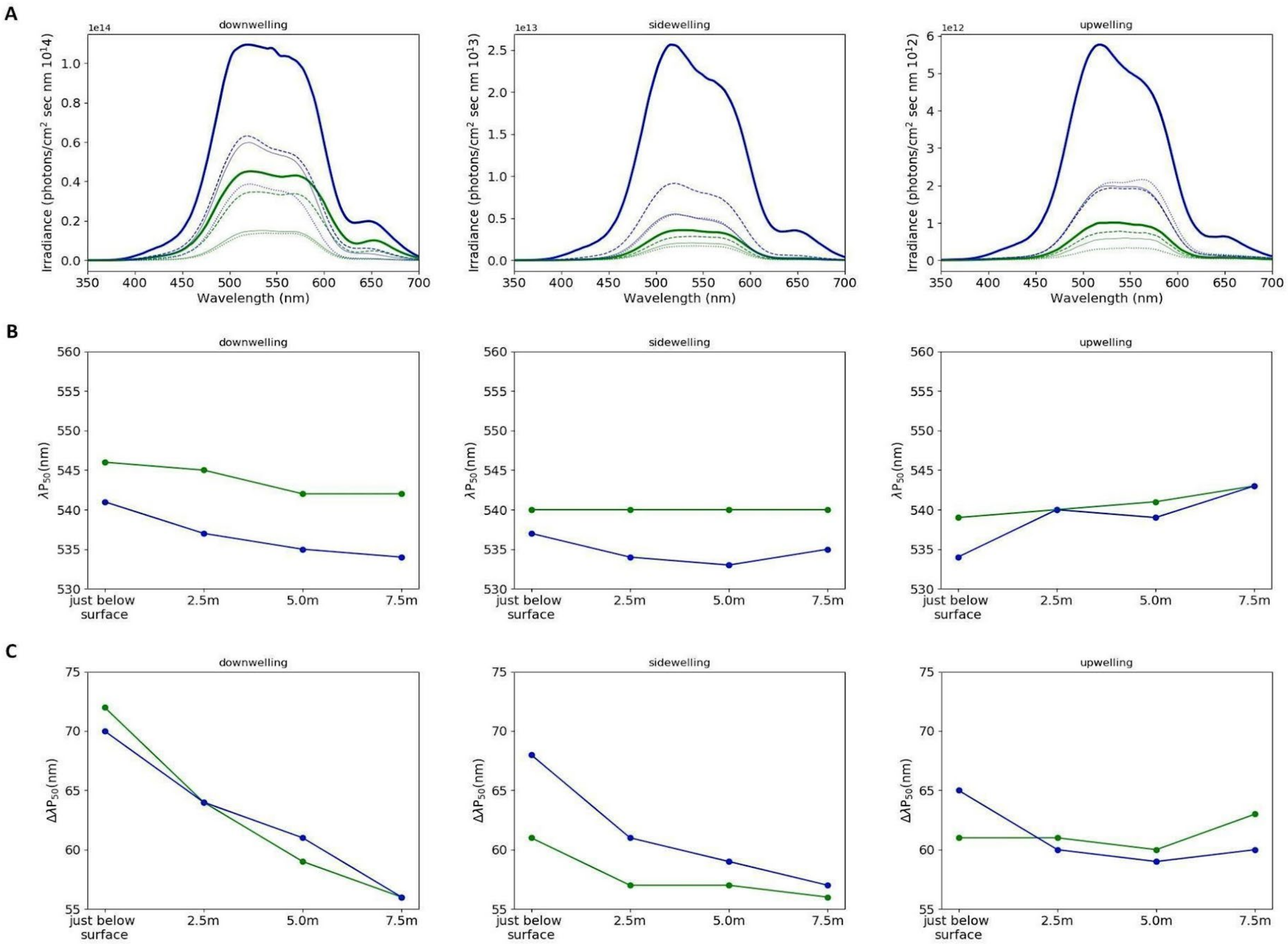

**Table S1.** Photon flux densities at different depths, obtained from in situ measurements at the collection sites, Curacao (CAR) and Isla Saboga (TEP), by integrating downwelling, sidewelling and upwelling spectral irradiances over the wavelength range 300-700nm.

| Direction of Light | Depth (m) | TEP | CAR |
|---|---|---|---|
| downwelling | under surface | 6,26725E+15 | 1,49655E+16 |
| | 2,5 | 4,3981E+15 | 7,48979E+15 |
| | 5 | 1,73959E+15 | 6,74466E+15 |
| | 7,5 | 1,53933E+15 | 4,10828E+15 |
| sidewelling | under surface | 4,197E+14 | 3,15857E+15 |
| | 2,5 | 3,17093E+14 | 1,03737E+15 |
| | 5 | 2,3279E+14 | 6,01511E+14 |
| | 7,5 | 1,90001E+14 | 5,87705E+14 |
| upwelling | under surface | 1,19826E+14 | 6,98744E+14 |
| | 2,5 | 9,1702E+13 | 2,26704E+14 |
| | 5 | 7,02535E+13 | 2,32913E+14 |
| | 7,5 | 4,13295E+13 | 2,51191E+14 |

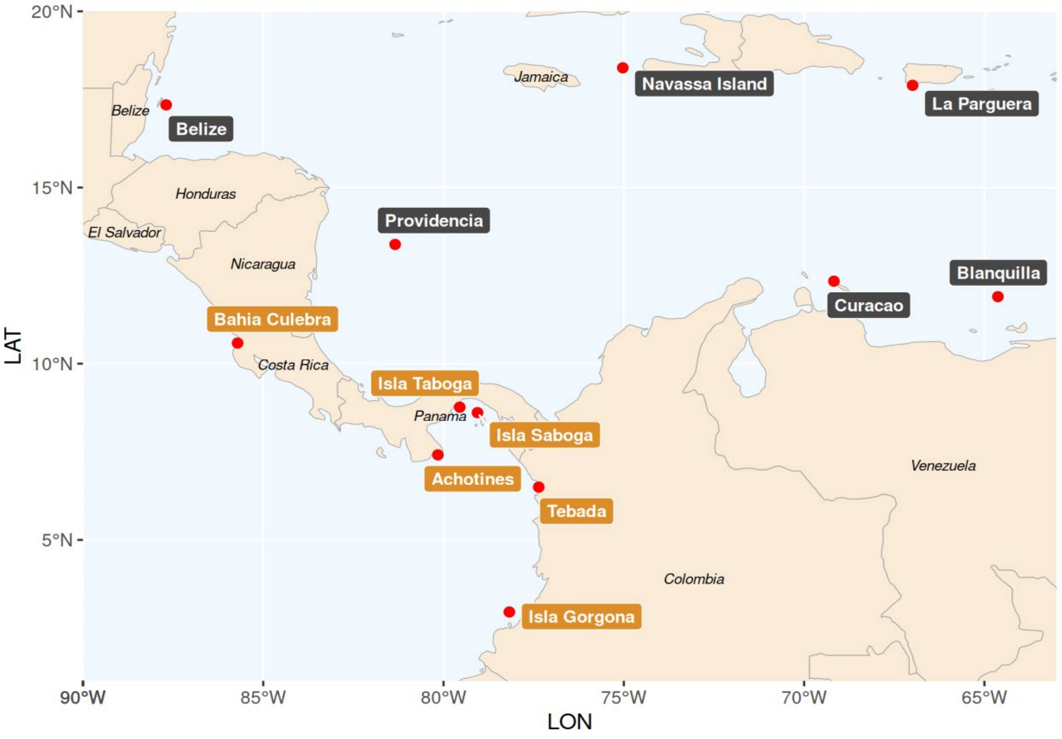


| Site | Oceanic Basin | LAT | LONG |
|---|---|---|---|
| Providencia (Colombia) | CAR | 13.385472 | -81.34161 |
| Kleine Knip (Curacao) | CAR | 12.341556 | -69.17378 |
| Blanquilla (Venezuela) | CAR | 11.902278 | -64.62647 |
| Turneffe Atoll (Belize) | CAR | 17.344983 | -87.69158 |
| La Parguera (Puerto Rico) | CAR | 17.900247 | -66.99238 |
| Navassa Island (USA) | CAR | 18.397714 | -75.02908 |
| Isla Saboga (Panama) | TEP | 8.6107722 | -79.05817 |
| Isla Taboga (Panama) | TEP | 8.765167 | -79.54992 |
| Achotines (Panama) | TEP | 7.4119056 | -80.15665 |
| Caleta Tebada (Colombia) | TEP | 6.4975417 | -77.36098 |
| Isla Gorgona, La Azufrada reef (Colombia) | TEP | 2.9533333 | -78.17491 |
| Bahia Culebra, Golfo de Papagayo (Costa Rica) | TEP | 10.585792 | -85.71094 |

**Figure S3.** Location and geographic coordinates of representative coral reef sites across the Caribbean Sea and the Tropical East Pacific from which the remote sensing reflectance data were extracted.

A.

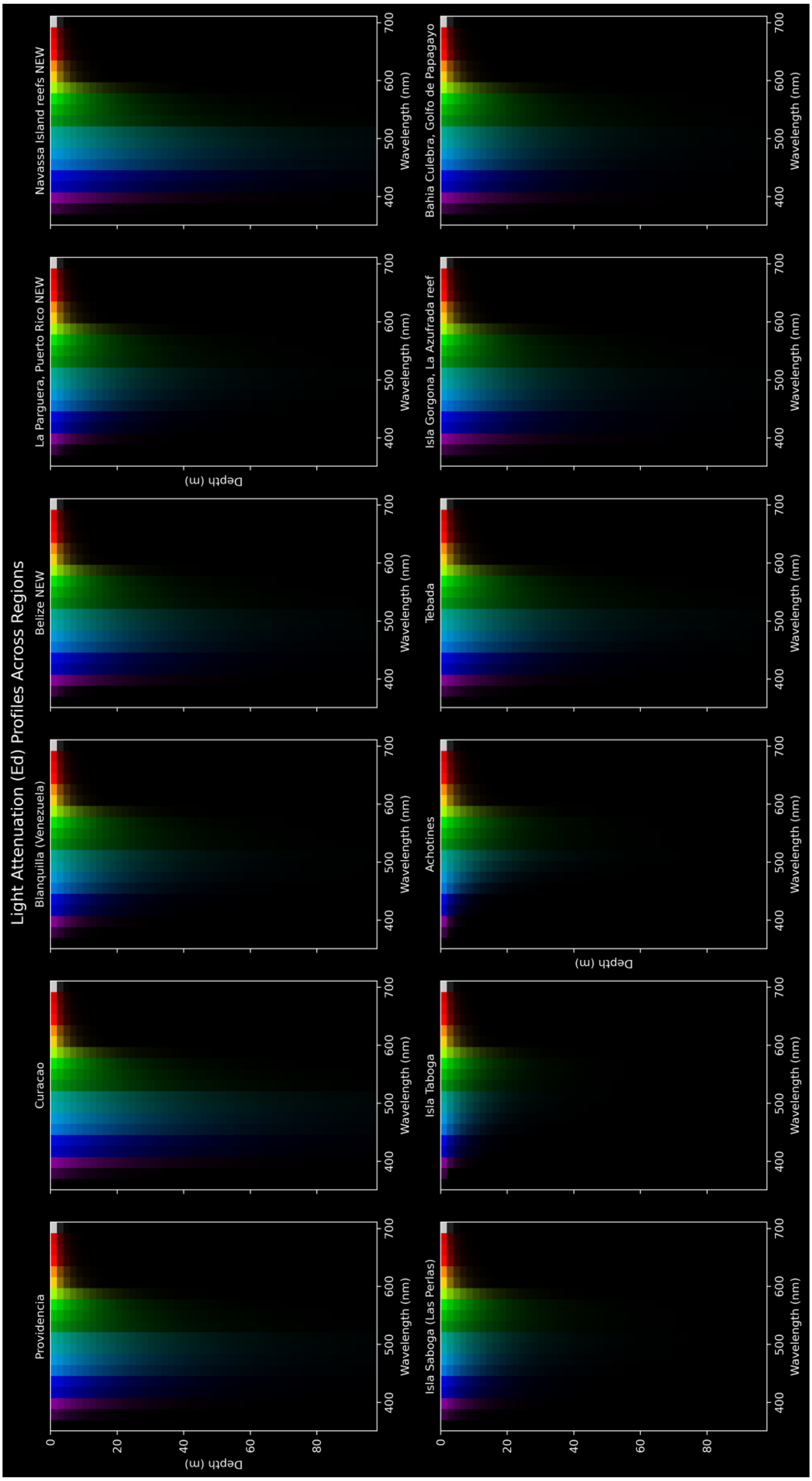
Light Attenuation (Ed) Profiles Across Regions
Providencia
Curacao
Blanquilla (Venezuela)
Belize NEW
La Parguera, Puerto Rico NEW
Navassa Island reefs NEW
Isla Saboga (Las Perlas)
Isla Taboga
Achotines
Tebada
Isla Gorgona, La Azufrada reef
Bahia Culebra, Golfo de Papagayo
Depth (m)
Wavelength (nm)
0
20
40
60
80
400
500
600
700

B.

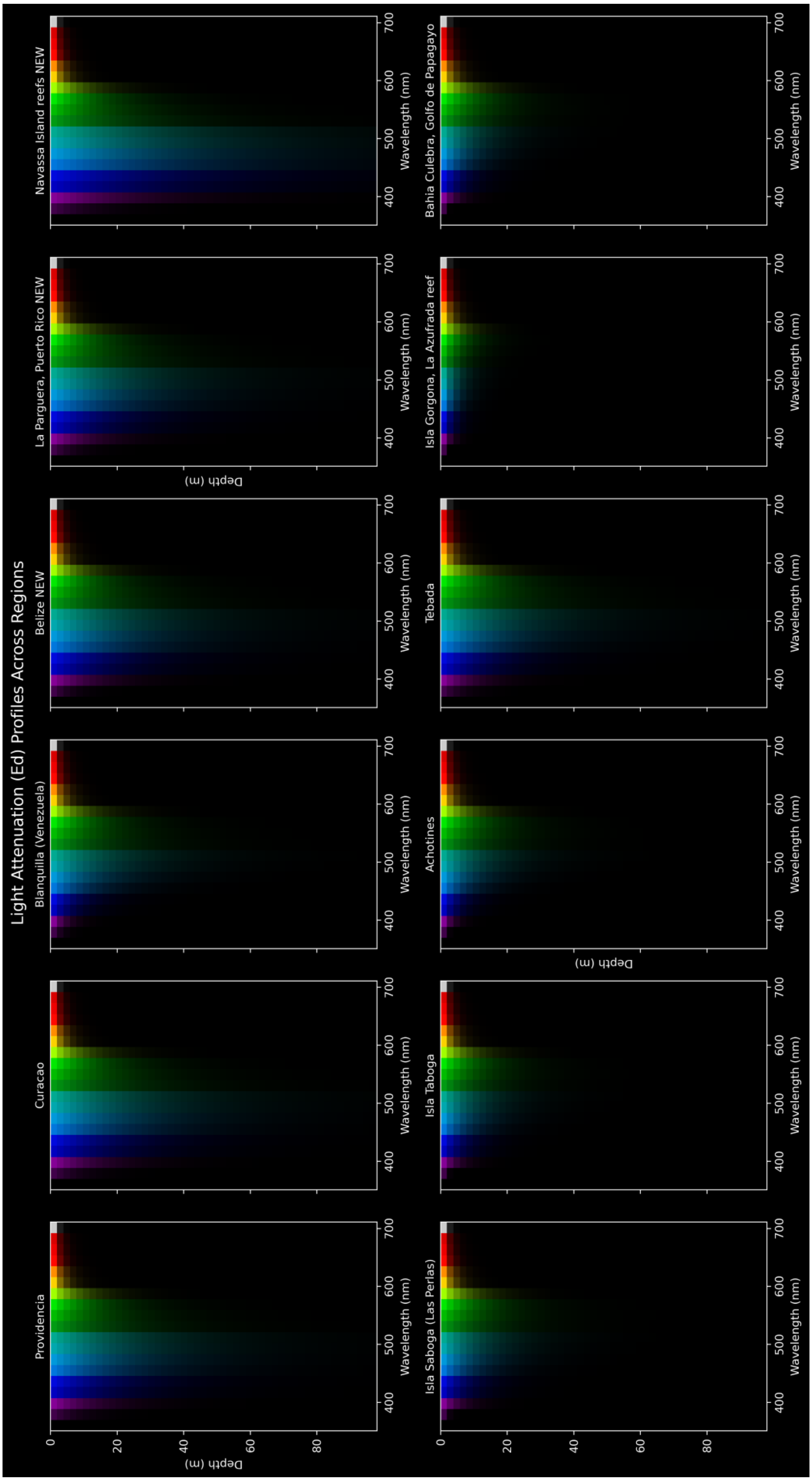
Light Attenuation (Ed) Profiles Across Regions
Providencia
Curacao
Blanquilla (Venezuela)
Belize NEW
La Parguera, Puerto Rico NEW
Navassa Island reefs NEW
Isla Saboga (Las Perlas)
Isla Taboga
Achotines
Tebada
Isla Gorgona, La Azufrada reef
Bahia Culebra, Golfo de Papagayo
Depth (m)
Wavelength (nm)

**Figure S4.** Approximation of the underwater light field at depth, across the twelve sites in the TEP and CAR, based on the remote sensing estimated irradiance profiles (A. September 2024, B. February 2025), assuming a water column of homogeneously distributed optical components; *upper row*, CAR sites; *lower row*, TEP sites.

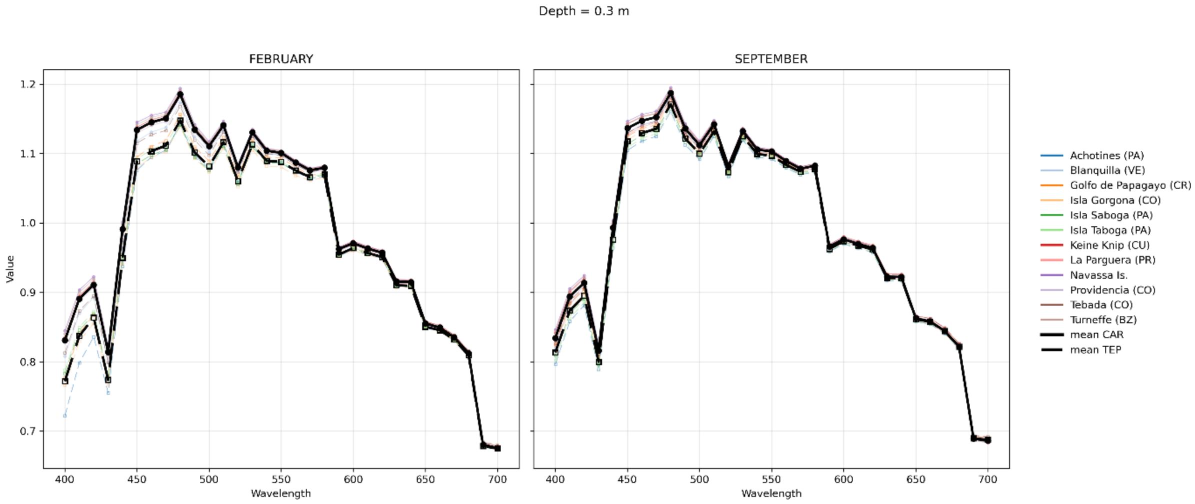
Depth = 0.3 m
FEBRUARY
SEPTEMBER
Value
Wavelength
Achotines (PA)
Blanquilla (VE)
Golfo de Papagayo (CR)
Isla Gorgona (CO)
Isla Saboga (PA)
Isla Taboga (PA)
Keine Knip (CU)
La Parguera (PR)
Navassa Is.
Providencia (CO)
Tebada (CO)
Turneffe (BZ)
mean CAR
mean TEP

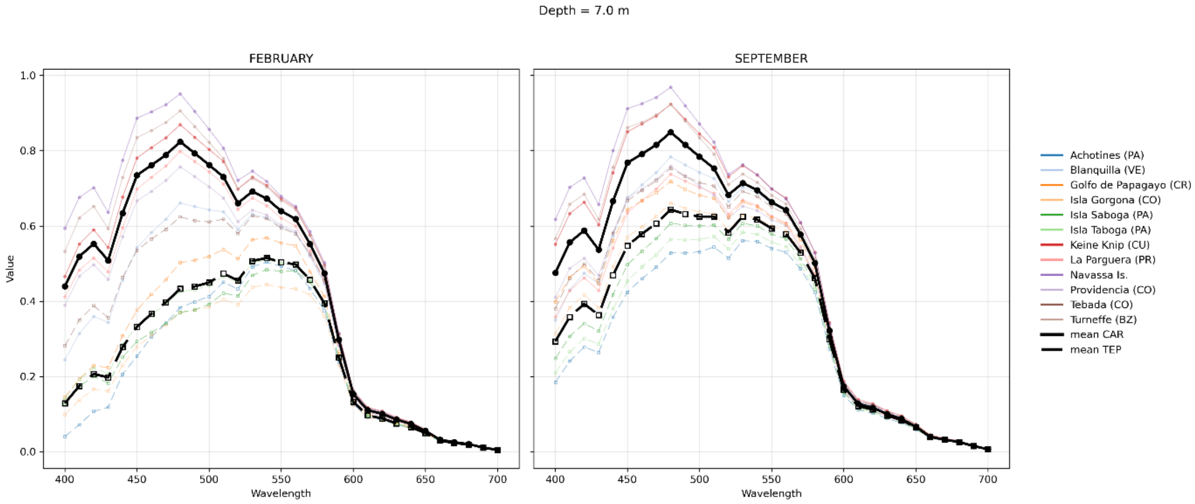
Depth = 7.0 m
FEBRUARY
SEPTEMBER
Value
Wavelength
Achotines (PA)
Blanquilla (VE)
Golfo de Papagayo (CR)
Isla Gorgona (CO)
Isla Saboga (PA)
Isla Taboga (PA)
Keine Knip (CU)
La Parguera (PR)
Navassa Is.
Providencia (CO)
Tebada (CO)
Turneffe (BZ)
mean CAR
mean TEP

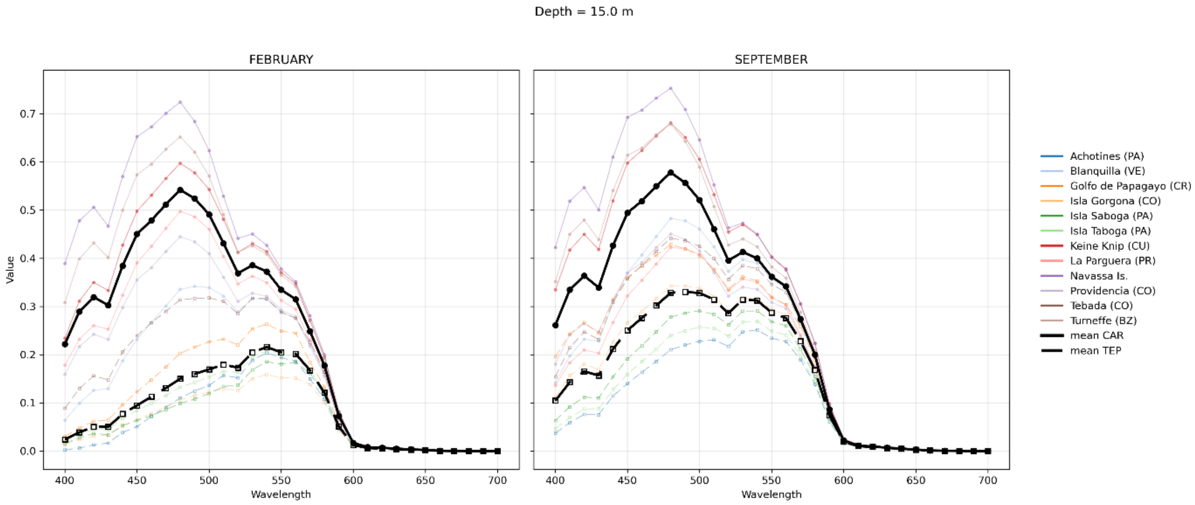
Depth = 15.0 m
FEBRUARY
SEPTEMBER
Value
Wavelength
Achotines (PA)
Blanquilla (VE)
Golfo de Papagayo (CR)
Isla Gorgona (CO)
Isla Saboga (PA)
Isla Taboga (PA)
Keine Knip (CU)
La Parguera (PR)
Navassa Is.
Providencia (CO)
Tebada (CO)
Turneffe (BZ)
mean CAR
mean TEP

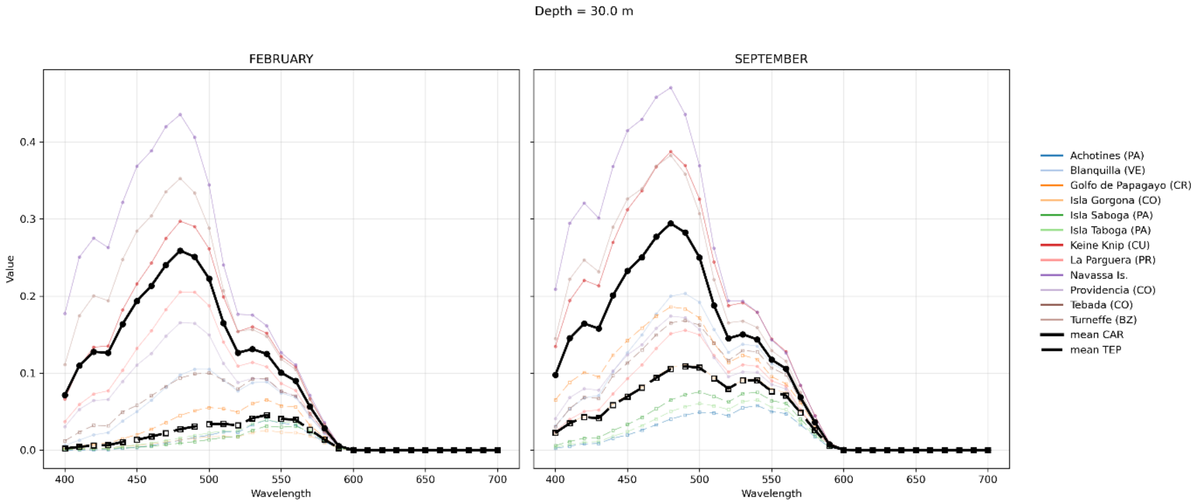


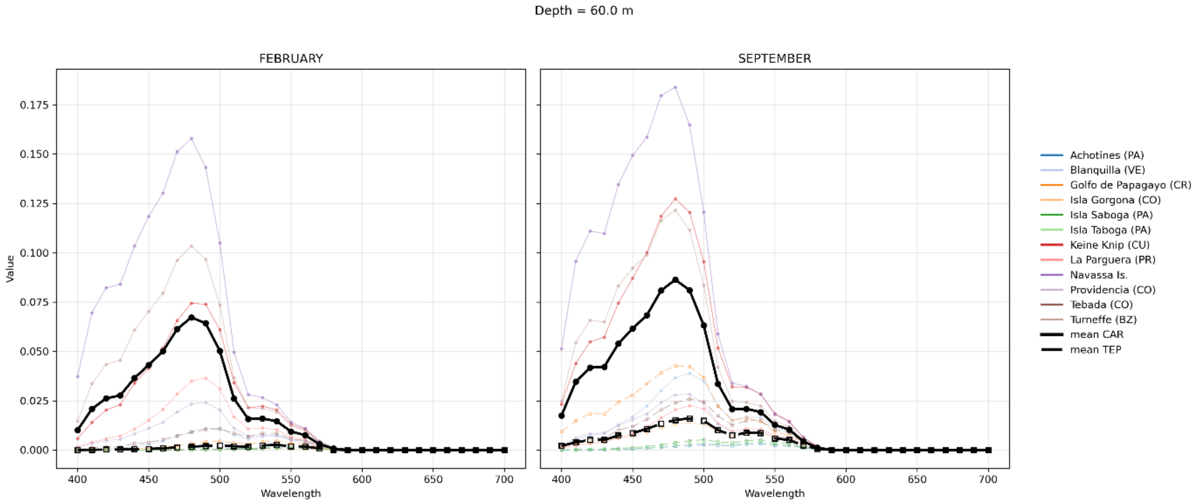


**Figure S5.** Remote sensing-derived median downwelling irradiances at different depths (0.3m, 7m, 15m, 30m, 60m), during (Feb) and outside (Sept) the upwelling season. Each line represents the median over 10 years, at each of twelve representative sites in the CAR (continuous lines) and TEP (dashed lines). Thicker lines indicate CAR and TEP averages across sites.

**A.**

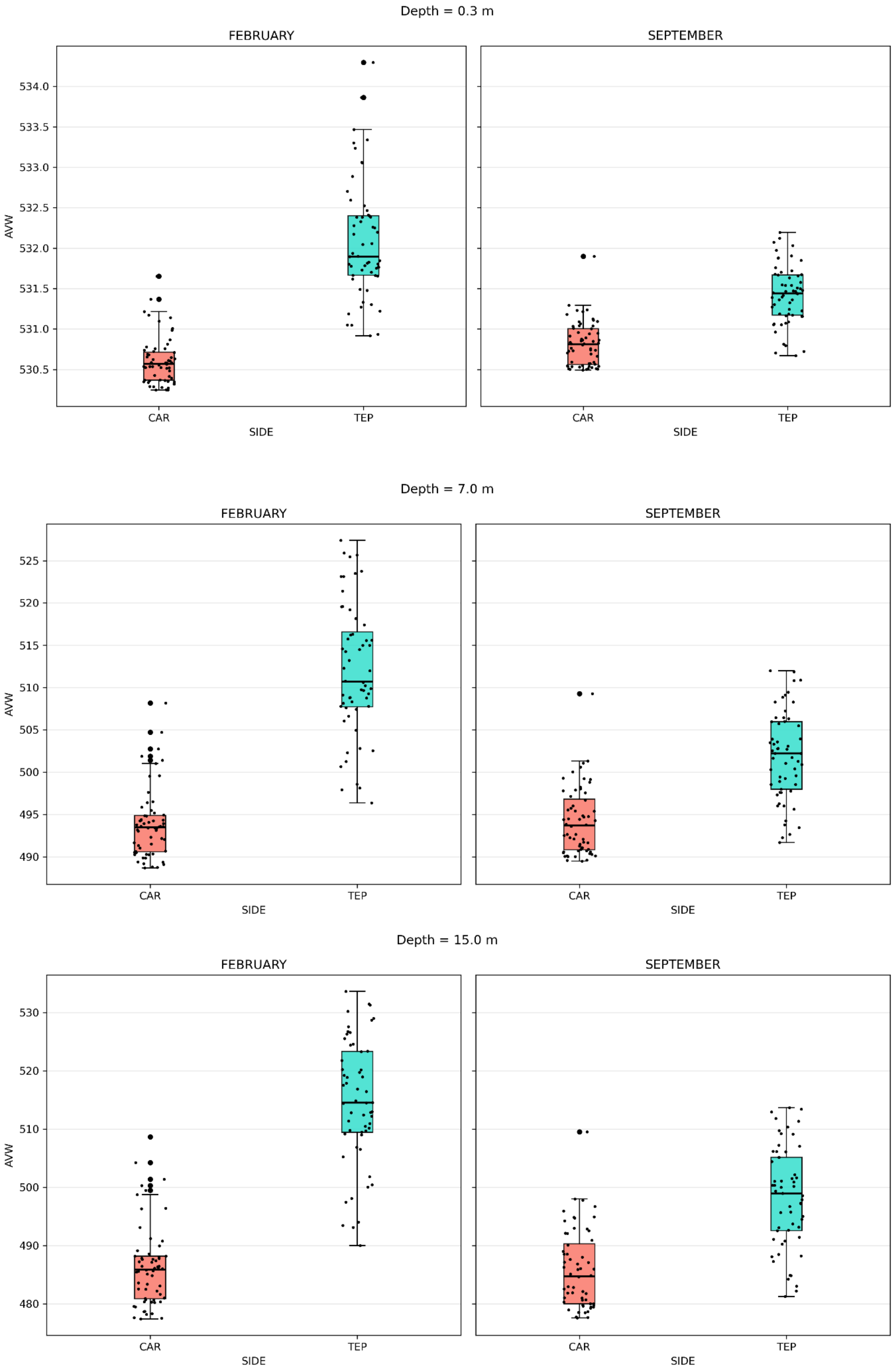
Depth = 0.3 m
FEBRUARY
SEPTEMBER
AVW
534.0
533.5
533.0
532.5
532.0
531.5
531.0
530.5
CAR
TEP
SIDE
Depth = 7.0 m
FEBRUARY
SEPTEMBER
AVW
525
520
515
510
505
500
495
490
CAR
TEP
SIDE
Depth = 15.0 m
FEBRUARY
SEPTEMBER
AVW
530
520
510
500
490
480
CAR
TEP
SIDE

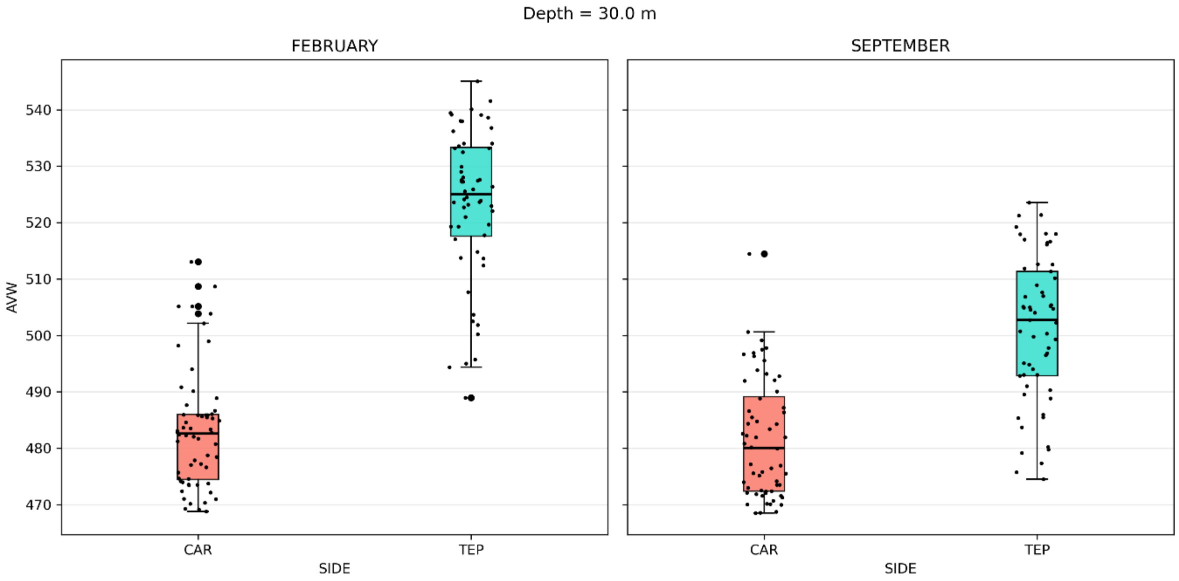
Depth = 30.0 m
FEBRUARY
SEPTEMBER
AVW
540
530
520
510
500
490
480
470
CAR
TEP
SIDE
CAR
TEP
SIDE

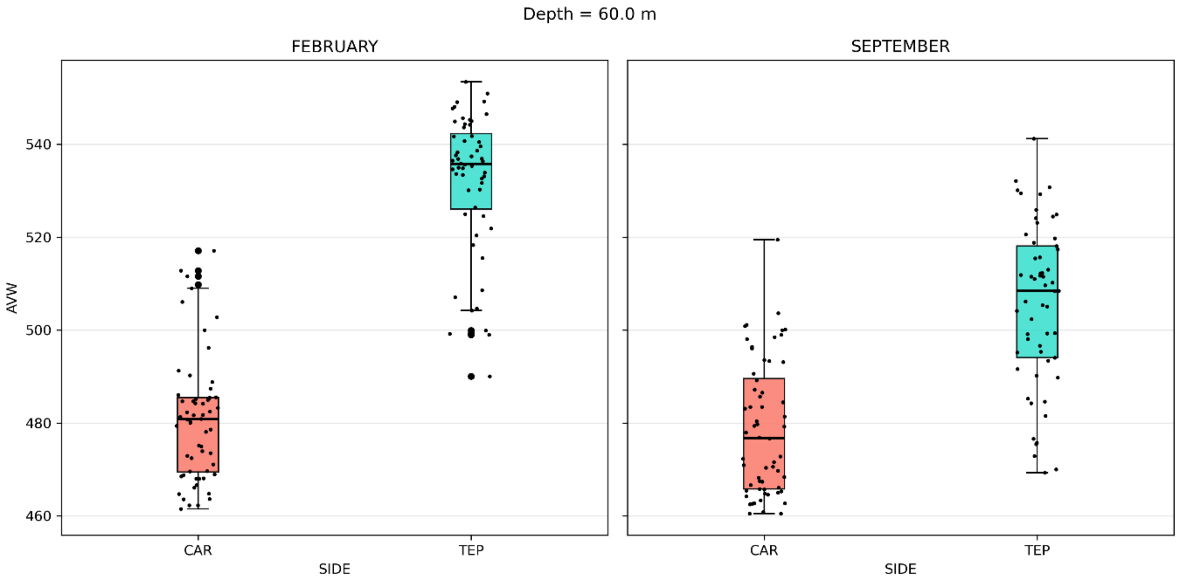
Depth = 60.0 m
FEBRUARY
SEPTEMBER
AVW
540
520
500
480
460
CAR
TEP
SIDE
CAR
TEP
SIDE

**B.**

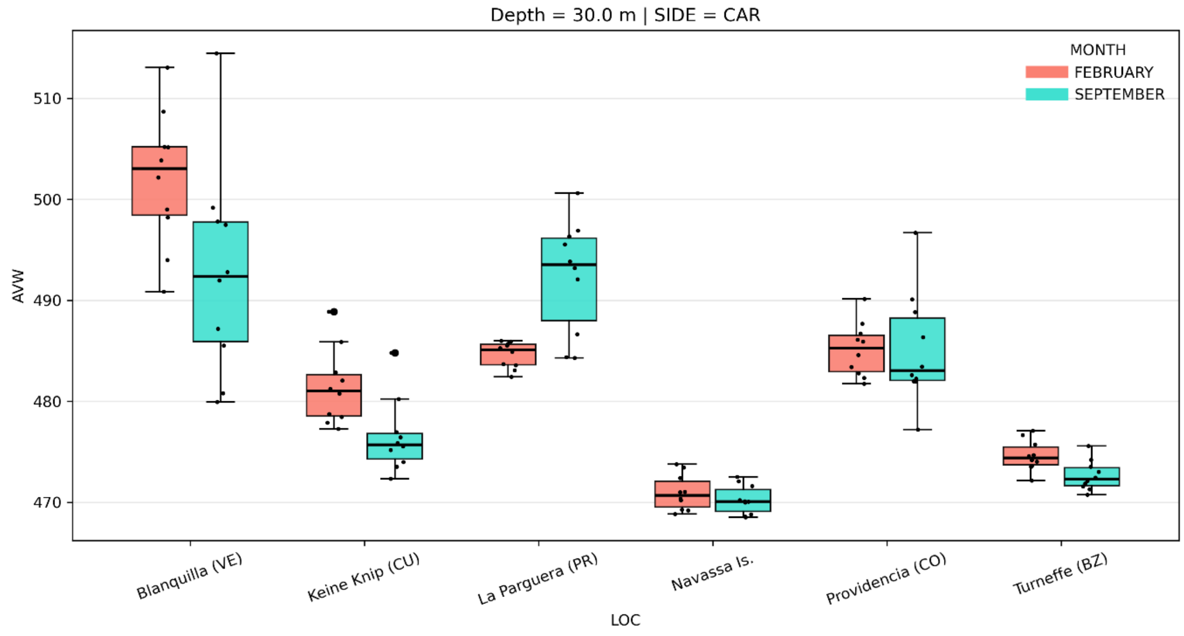


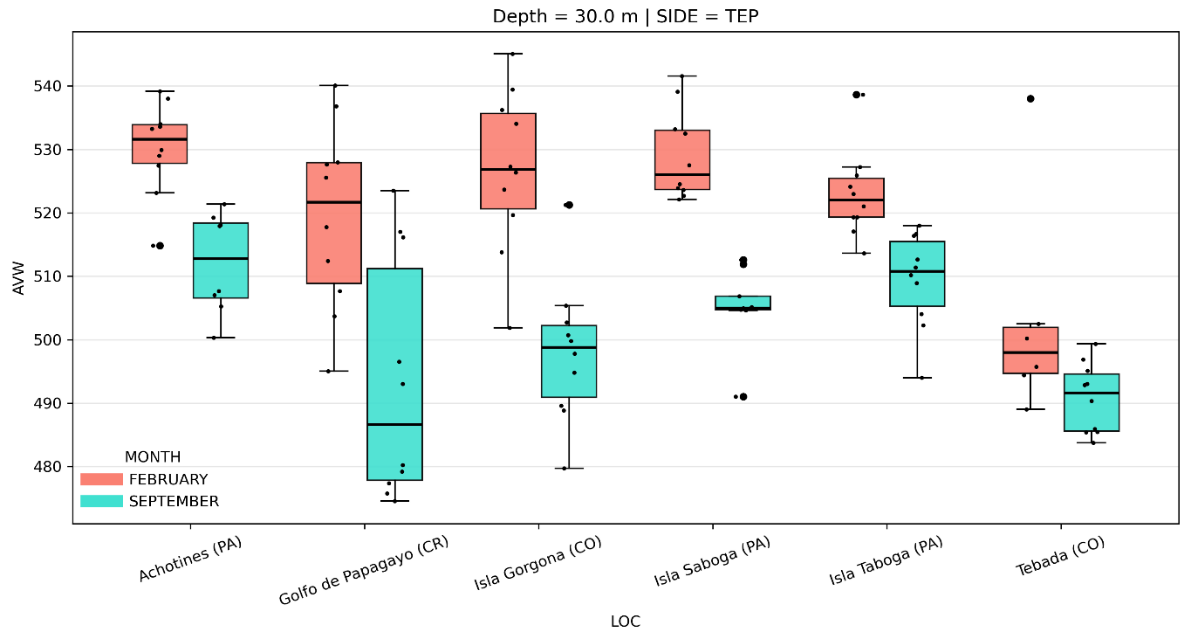


**Figure S6.** A. The Apparent Visible Wavelength (AVW) at different depths (0.3m, 7m, 15m, 30m, 60m), pooled by oceanic basin, during (Feb) and outside (Sept) the upwelling seasons; B. the same, by individual locality, at 30m: *above*, CAR sites; *below*, TEP sites.

**Table S2.** GenBank accession numbers for the visual opsin gene sequences from a range of teleost species used in the opsin gene trees (Figs. S7-S11).

| Opsin Gene Tree | Opsin Gene Accession Numbers |
|---|---|
| RH1 | *Danio rerio* RH1: AB087811.1, *Takifugu rubripes* RH1: AF201471.1, *Epinephelus fuscoguttatus* RH1: AP022681.1, *Epinephelus bruneus* RH1: LC064406.1, *Epinephelus moara* RH1: XM_050065809.1, *Maylandia/Metraclima zebra* RH1: AY775114.1, *Abudefduf sexfasciatus* RH1: HQ286548.1, *Dascyllus trimaculatus* RH1: HQ286552.1, *Chromis viridis* RH1: HQ286550.1, *Oryzias latipes* RH1: AB180742.1 |
| RH2 | *Danio rerio* RH2-3: NC_007117.6; RH2-1: NC_007117.6; RH2-2: NC_007117.6, *Paralichthys olivaceus* RH2B1: MN400438.1, RH2A1: MN400436.1 *Cheilodipterus artus* RH2B2 MH991738.1, *Maylandia/Metraclima zebra* RH2A: JF262089.1, RH2A beta: XM_004548752.5, RH2B: JF262089.1, *Oryzias latipes* RH2B: AB223054.1, RH2A: AB223053.1, *Abudefduf sexfasciatus* RH2A: HQ286528.1, RH2B: HQ286518.1, *Chromis viridis* RH2A: HQ286530.1, RH2B: HQ286520.1, *Dascyllus trimaculatus* RH2A: HQ286532.1, RH2B: HQ286522.1, *Plectropomus leopardus* RH2B: XM_042507605.1, RH2A1: XM_042507581.1, RH2A2: XM_042507592.1, *Epinephelus fuscoguttatus* RH2A1: XM_049584805.1, RH2B: XM_049584813.1, *Epinephelus moara* RH2A1: XM_050038009.1, RH2A2: XM_050038011.1, RH2B: XM_050038010.1, *Epinephelus lanceolatus* RH2A1: XM_033627346.2, RH2A2: XM_033627347.2, *Chelmon rostratus* RH2A2: XM_041946754.1, RH2A1 alpha: XM_041946743.1, RH2A1 beta: XM_041954010.1, *Scophthalmus maximus* RH2A2: XM_035632464.1, *Takifugu rubripes* RH2: AF226989.1, *Euthynnus affinis* Rh2A1: LC016737.1, *Sebastes umbrosus* RH2A1 alpha: XM_037768925.1, RH2A1 beta XM_037775992.1, *Epinephelus bruneus* RH2A1: LC064408.1, RH2A2: LC064411.1, *Ostorhinchus angustatus* RH2A1: MH979515.1, *Aulonocara baenschi* RH2A beta: GQ422500.1 |
| LWS | *Danio rerio* LWS2: NM_001002443.2, LWS1: NM_001313715.1, *Maylandia/Metraclima zebra* LWS: AF247126.1, *Epinephelus moara* LWS: XM_050037641.1, *Epinephelus bruneus* LWS: LC064414.1, *Ostorhinchus angustatus* LWS: MH979542.1, *Takifugu rubripes* LWS: XM_003973673.3, *Abudefduf sexfasciatus* LWS: HQ286538.1, *Dascyllus trimaculatus* LWS: HQ286546.1, *Chromis viridis* LWS: HQ286545.1, *Oryzias latipes* LWS-A:AB223056.1, LWS-B: AB223052.1 |
| SWS1 | *Danio rerio* SWS1: KT008399.1, *Epinephelus bruneus* SWS1: LC016745.1, *Epinephelus moara* SWS1: XM_050036791.1, *Epinephelus fuscoguttatus* SWS1:XM_049567370.1, *Abudefduf sexfasciatus* SWS1: HQ286498.1, *Dascyllus trimaculatus* SWS1: HQ286502.1, *Chromis viridis* SWS1: HQ286500.1, *Oryzias latipes* SWS1: AB223058.1, *Maylandia/Metraclima* |

| | |
|---|---|
| | *zebra* SWS1:JF262085.1 |
| SWS2 | Danio rerio SWS2: KT008398.1, *Oryzias latipes* SWS2B: AB223057.1, SWS2A: AB223056.1, Ostorhinchus angustatus SWS2A alpha: KP004341.1 , SWS2A beta: KP004345.1, *Epinephelus fuscoguttatus* SWS2B: XM_049577827.1, SWS2A2: XM_049577820.1, SWS2A1: XM_049577841.1, *Epinephelus bruneus* SWS2A2: LC064413.1, SWS2A1: LC064412.1, *Epinephelus moara* SWS2B: XM_050038296.1, SWS2A2: XM_050038295.1, SWS2A1: XM_050038297.1, *Maylandia/Metraclima zebra* SWS2A: AF247114.1, SWS2B: AF247118.1, *Abudefduf sexfasciatus* SWS2B: HQ286508.1, *Dascyllus trimaculatus* SWS2B: HQ286512.1, *Chromis viridis* SWS2B: HQ286510.1 |

135

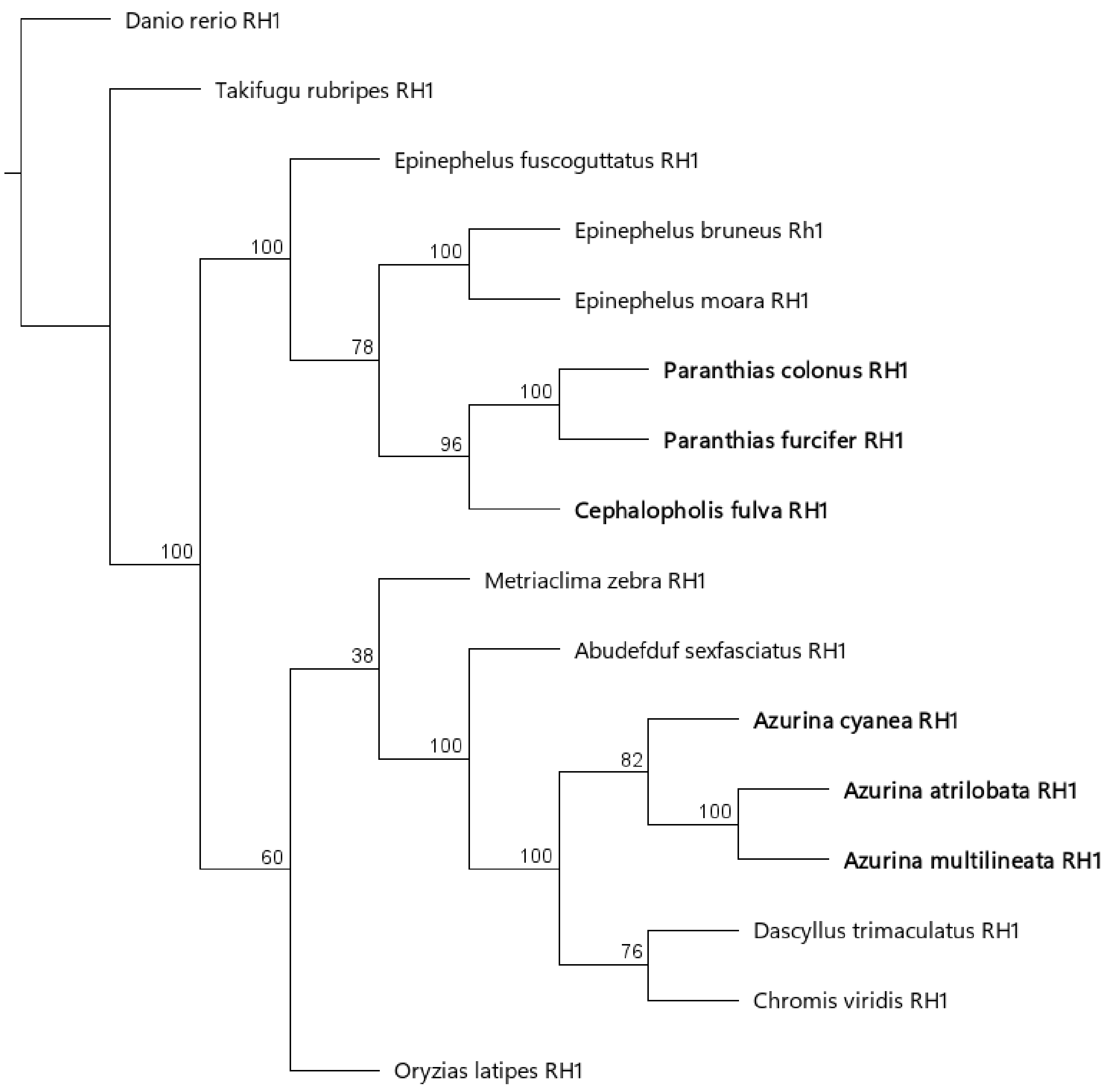

**Figure S7.** Maximum likelihood phylogenetic tree of RH1 opsin gene sequences from a range of teleost species obtained from GenBank, grouper and damselfish sequences recovered in this study are shown in bold. The bootstrap percent confidence values are shown for each branch. The *Danio rerio* RH1 sequence was used to root the tree (GenBank accession No.: AB087811.1). Accession numbers for all other sequences used are listed in Supplementary Table S3.

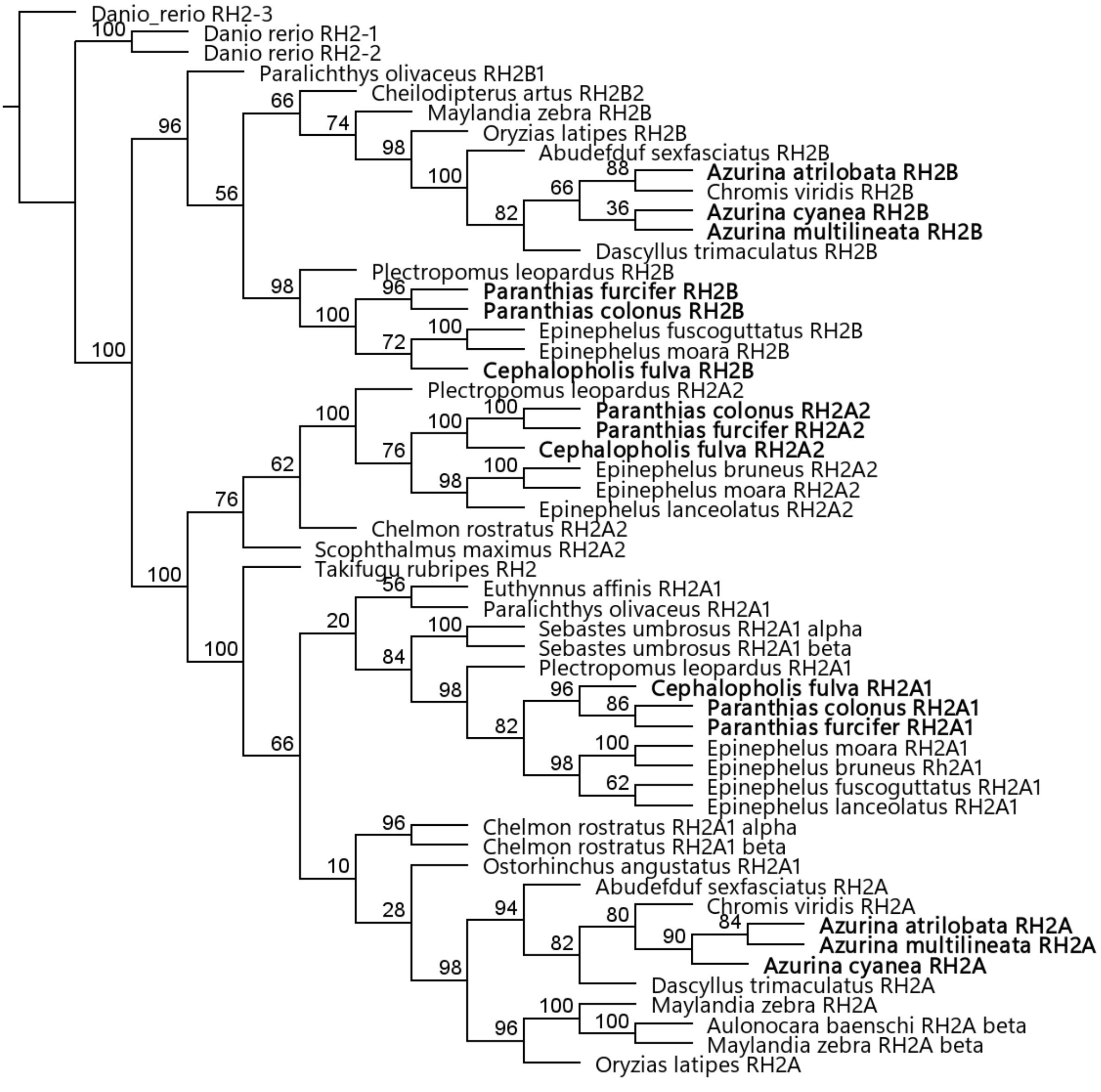

**Figure S8.** Maximum likelihood phylogenetic tree of RH2 opsin gene sequences from a range of teleost species obtained from GenBank, grouper and damselfish sequences recovered in this study are shown in bold. The bootstrap percent confidence values are shown for each branch. The *Danio rerio* RH2-3 sequence was used to root the tree (GenBank accession No.: NC_007117.6). Accession numbers for all other sequences used are listed in Supplementary Table S3.

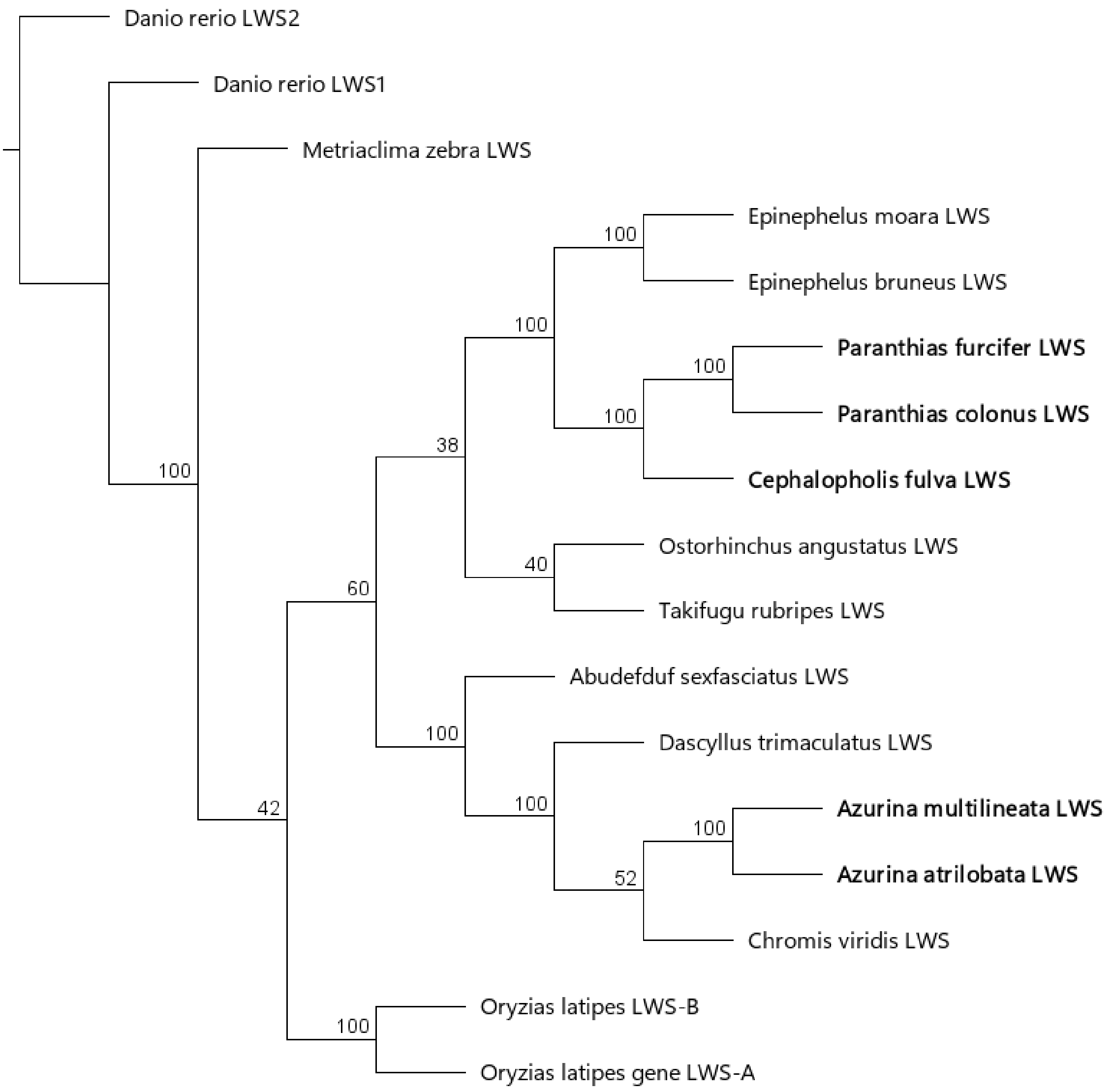

**Figure S9.** Maximum likelihood phylogenetic tree of LWS opsin gene sequences from a range of teleost species obtained from GenBank, grouper and damselfish sequences recovered in this study are shown in bold. The bootstrap percent confidence values are shown for each branch. The *Danio rerio* LWS2 sequence was used to root the tree (GenBank accession No.: NM_001002443.2). Accession numbers for all other sequences used are listed in Supplementary Table S3.

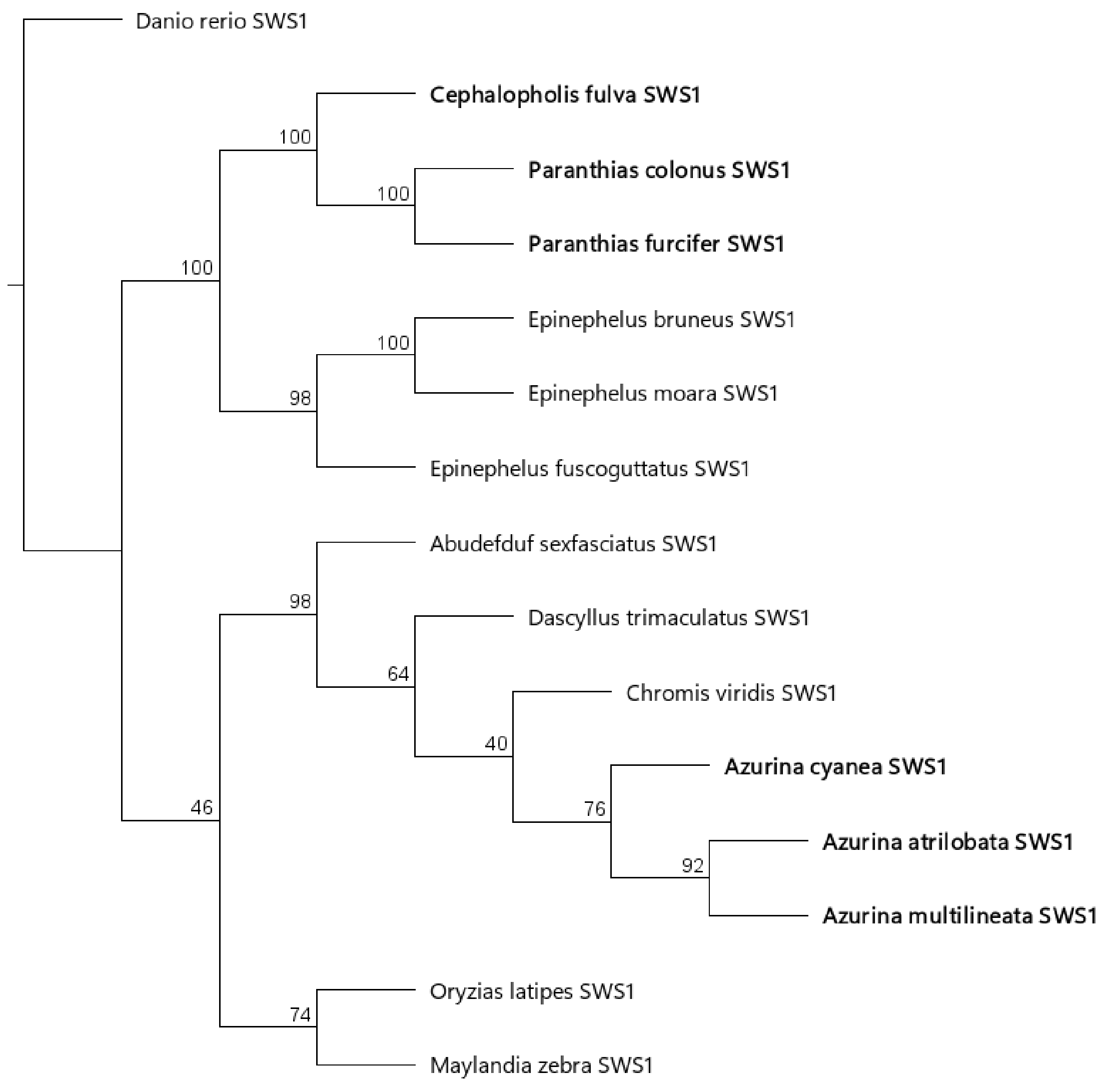

**Figure S10.** Maximum likelihood phylogenetic tree of SWS1 opsin gene sequences from a range of teleost species obtained from GenBank, grouper and damselfish sequences recovered in this study are shown in bold. The bootstrap percent confidence values are shown for each branch. The *Danio rerio* SWS1 sequence was used to root the tree (GenBank accession No.: KT008399.1). Accession numbers for all other sequences used are listed in Supplementary Table S3.

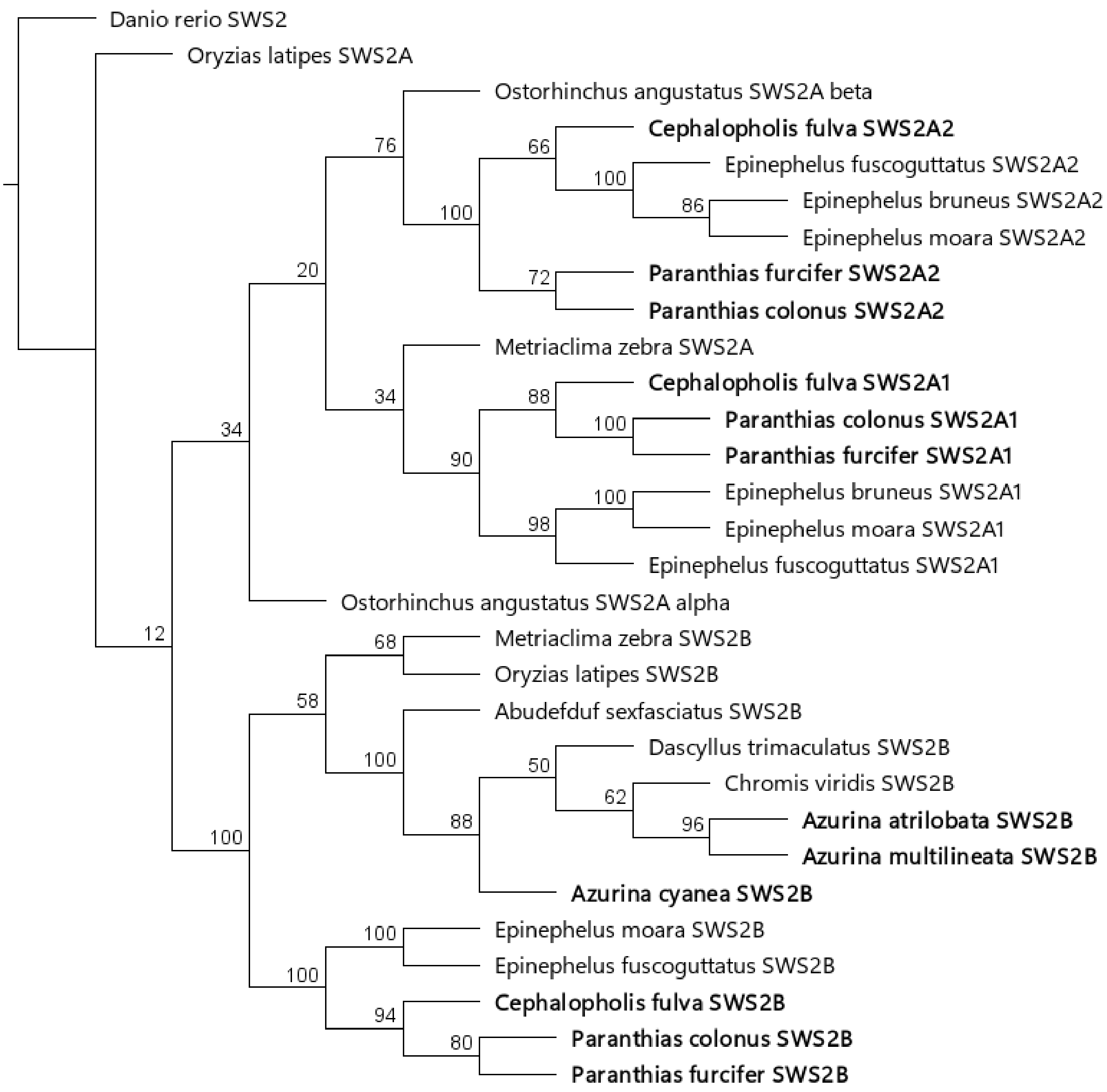


**Figure S11.** Maximum likelihood phylogenetic tree of SWS2 opsin gene sequences from a range of teleost species obtained from GenBank, grouper and damselfish sequences recovered in this study are shown in bold. The bootstrap percent confidence values are shown for each branch. The *Danio rerio* SWS2 sequence was used to root the tree (GenBank accession No.: KT008398.1). Accession numbers for all other sequences used are listed in Supplementary Table S3.

**Table S3.** Wavelength of maximum sensitivity ($\lambda_{max}$) of visual pigments measured in photoreceptors in situ with micro-spectrophotometry and proposed relationship with expressed opsin genes.

| | Rod | Short Wave Sensitive | | | | Medium Wave Sensitive | | | | | | Long Wave Sensitive |
|---|---|---|---|---|---|---|---|---|---|---|---|---|
| | RH1 | SWS1 | SWS2B | SWS2A1 | SWS2A2 | RH2B | RH2B+RH2A | RH2A+RH2B | RH2A1 | RH2A2 | RH2A | LWS |
| ***Pomacentridae*** | | | | | | | | | | | | |
| *Azurina atrilobata* | 495.98 (1.14) | - | 412.0 (1.63) | - | - | 476.0 (2.0) | 485.59 (1.54) | 506.2 (1.1) | - | - | 515.67 (1.15) | 537 |
| | n=45 | - | n=4 | - | - | n=6 | n=17 | n=5 | - | - | n=3 | n=1 |
| *Azurina multilineata* | 495.66 (1.38) | 361 | 408.67 (0.82) | - | - | 475.0 (2.27) | 483.89 (2.26) | 506.6 (2.07) | - | - | 517.89 (1.05) | 534.93 (1.91) |
| | n=64 | n=1 | n=6 | - | - | n=47 | n=9 | n=5 | - | - | n=9 | n=15 |
| *Azurina cyanea* | 490.13 (2.31) | - | 403 | - | - | 477.96 (3.47) | - | 507 | - | - | - | 529.0 (1.41) |
| | n=24 | - | n=1 | - | - | n=55 | - | n=1 | - | - | - | n=2 |
| ***Serranidae*** | | | | | | | | | | | | |
| *Paranthias colonus* | 496.59 (1.21) | - | - | 444.0 (1.22) | 454.89 (1.97) | - | - | - | 507.55 (1.63) | 517.7 (1.15) | - | 527.88 (2.47) |
| | n=39 | - | - | n=5 | n=18 | - | - | - | n=11 | n=46 | - | n=16 |
| *Paranthias furcifer* | 483.27 (1.74) | - | - | 434.5 (0.71) | 443.5 (2.12) | - | - | - | - | - | - | 529.5 (2.12) |
| | n=11 | - | - | n=2 | n=2 | - | - | - | - | - | - | n=2 |
| *Cephalopholis fulva* | 495.94 (1.92) | - | - | 433.0 (1.15) | 454.33 (3.51) | - | - | - | 507.33 (3.01) | 518.2 (0.45) | - | 529.27 (1.94) |
| | n=18 | - | - | n=4 | n=3 | - | - | - | n=6 | n=5 | - | n=37 |

**Table S4.** Summary of variable sites in opsin gene sequences recovered in the damselfish and the grouper geminate contrasts. Location of tuning sites, retinal binding pockets and transmembrane regions were taken from Hofmann et al. (2012).

| Species Set | Opsin | Number of Variable Sites | | | |
|---|---|---|---|---|---|
| | | *Known tuning site* | *Retinal binding pocket* | *Transmembrane region* | *Full region* |
| *Damselfish* | *RH1* | 0 | 0 | 4 | 5 |
| *Damselfish* | *SWS1* | 0 | 1 | 5 | 6 |
| *Damselfish* | *SWS2B* | 1 | 1 | 8 | 14 |
| *Damselfish* | *RH2B* | 1 | 2 | 12 | 14 |
| *Damselfish* | *RH2A* | 0 | 1 | 7 | 9 |
| *Damselfish* | *LWS* | 1 | 1 | 9 | 14 |
| *Grouper* | *RH1* | 2 | 2 | 4 | 4 |
| *Grouper* | *SWS1* | 0 | 0 | 1 | 1 |
| *Grouper* | *SWS2B* | 0 | 0 | 0 | 0 |
| *Grouper* | *SWS2A1* | 1 | 1 | 3 | 4 |
| *Grouper* | *SWS2A2* | 0 | 1 | 1 | 1 |
| *Grouper* | *RH2B* | 0 | 1 | 4 | 5 |
| *Grouper* | *RH2A1* | 0 | 0 | 3 | 3 |
| *Grouper* | *RH2A2* | 0 | 0 | 3 | 4 |
| *Grouper* | *LWS* | 0 | 0 | 1 | 3 |

**Table S5.** Known amino acid tuning sites recovered in the damselfish contrast and their estimated effects. Amino acid site number is indicated and in brackets, the corresponding site in the bovine rhodopsin reference. Cases where estimates of the size of a given tuning shift could not be found are indicated with a question mark. MSP estimates from other species used as reference were taken from the following: Stieb et al. (2016): *A. amboinensis*, Parry et al. (2005): *M. zebra*, Matsumoto et al. (2006); *O. latipes*, Escobar-Camacho et al. (2020): *R. guatemalensis,* Spady et al. (2006): *O. niloticus.*

| **Species** | **Variable sites** | | | | **Tuning effect (nm)** | | | | **Estimated $\lambda_{max}$ (nm)** |
|---|---|---|---|---|---|---|---|---|---|
| **RH2B** | 208 (207) | 125 (124) | - | - | 208 (207) | 125 (124) | - | - | |
| *A. atrilobata* | M | A | - | - | 0 | -2 | - | - | 478 |
| *A. multilineata* | L | S | - | - | -7 | 0 | - | - | 473 |
| *A. cyanea* | L | S | - | - | -7 | 0 | - | - | 473 |
| *P. amboinensis* | M | S | - | - | - | - | - | - | - |
| **SWS2B** | 49 (43) | 124 (118) | 174 (168) | 271 (265) | 49 (43) | 124 (118) | 174 (168) | 271 (265) | |
| *A. atrilobata* | F | T | S | Y | 0 | 0 | 4 | -15 | 412 |
| *A. multilineata* | L | T | S | Y | -3 | 0 | 4 | -15 | 409 |
| *A. cyanea* | F | S | S | Y | 0 | -10 | 4 | -15 | 402 |
| *M. zebra* | F | T | A | W | - | - | - | - | - |
| **RH1** | 83 (83) | 292 (292) | 299 (299) | - | 83 (83) | 292 (292) | 299 (299) | - | |
| *A. atrilobata* | N | A | S | - | -6 | - | - | - | 496 |
| *A. multilineata* | N | A | S | - | -6 | - | - | - | 496 |

| | | | | | | | | | |
|---|---|---|---|---|---|---|---|---|---|
| *A. cyanea* | N | S | A | - | -6 | -10 | 0 | - | 486 |
| *O. latipes* | D | A | S | - | - | - | - | - | - |
| **LWS** | 282 (269) | - | - | - | 282 (269) | - | - | - | |
| *A. atrilobata* | A | - | - | - | -16 | - | - | - | 514 |
| *A. multilineata* | T | - | - | - | - | - | - | - | 530 |
| *A. cyanea* | - | - | - | - | - | - | - | - | - |
| *R. guatemalensis* | T | - | - | - | - | - | - | - | - |
| **SWS1** | 90 (97) | - | - | - | 90 (97) | - | - | - | |
| *A. atrilobata* | S | - | - | - | ? | - | - | - | 360+/- tuning effect |
| *A. multilineata* | S | - | - | - | ? | - | - | - | 360+/- tuning effect |
| *A. cyanea* | A | - | - | - | 0 | - | - | - | 360 |
| *O. niloticus* | A | - | - | - | - | - | - | - | - |
| **RH2A** | - | - | - | - | - | - | - | - | |
| *A. atrilobata* | - | - | - | - | - | - | - | - | 518 |
| *A. multilineata* | - | - | - | - | - | - | - | - | 518 |
| *A. cyanea* | - | - | - | - | - | - | - | - | 518 |
| *O. niloticus* | - | - | - | - | - | - | - | - | - |

**Table S6.** Known amino acid tuning sites recovered in the grouper contrast and their estimated effects. Amino acid site number is indicated and in brackets, the corresponding site in the bovine rhodopsin reference. Cases where estimates of the size of a given tuning shift could not be found or where there was not enough information to provide a $\lambda_{max}$ estimate for the sequence are indicated with a question mark. Matsumoto et al. (2006); *O. latipes*, Steib et al. (2016): *A. amboinensis*, Cortesi et al. (2015): *P. fuscus,* Parry et al. (2005): *M. zebra*, Spady et al. (2006): *O. niloticus,* Escobar-Camacho et al. (2020): *R. guatemalensis.*

| **Species** | **Variable sites** | | | **Tuning effect (nm)** | | | **Estimated $\lambda_{max}$ (nm)** |
|---|---|---|---|---|---|---|---|
| **RH1** | 83 (83) | 292 (292) | 299 (299) | 83 (83) | 292 (292) | 299 (299) | |
| *C. colonus* | N | A | S | -6 | - | - | 496 |
| *C. furcifer* | N | S | A | -6 | -10 | 0 | 486 |
| *C. fulva* | N | A | A | -6 | - | 0 | 496 |
| *O. latipes* | D | A | S | - | - | - | - |
| **SWS1** | 45 (52) | - | - | 45 (52) | - | - | |
| *C. colonus* | T | - | - | - | - | - | 370 |
| *C. furcifer* | T | - | - | - | - | - | 370 |
| *C. fulva* | M | - | - | ? | - | - | 370+/- tuning effect |
| *P. amboinensis* | T | - | - | - | - | - | - |
| **SWS2A1** | 122 (116) | 305 (299) | - | 122 (116) | 305 (299) | - | |
| *C. colonus* | M | T | - | - | 2 | - | 450 |

| | | | | | | | |
|---|---|---|---|---|---|---|---|
| *C. furcifer* | T | T | - | ? | 2 | - | 450+/- tuning effect |
| *C. fulva* | M | T | - | - | 2 | - | 450 |
| *P. fuscus* | M | A | - | - | - | - | - |
| **SWS2A2** | 222 (216) | 275 (269) | 305 (299) | 222 (216) | 275 (269) | 305 (299) | |
| *C. colonus* | F | A | T | -8 | 0 | 2 | 451 |
| *C. furcifer* | F | A | T | -8 | 0 | 2 | 451 |
| *C. fulva* | F | T | T | -8 | - | 2 | 451 |
| *P. fuscus* | L | T | A | - | - | - | - |
| **SWS2B** | | - | - | - | - | - | |
| *C. colonus* | - | - | - | - | - | - | 423 |
| *C. furcifer* | - | - | - | - | - | - | 423 |
| *C. fulva* | - | - | - | - | - | - | 423 |
| *M. zebra* | - | - | - | - | - | - | - |
| **RH2A1** | 166 (158) | - | - | 166 (158) | - | - | |
| *C. colonus* | L | - | - | -10 | - | - | 518 |
| *C. furcifer* | L | - | - | -10 | - | - | 518 |
| *C. fulva* | L | - | - | -10 | - | - | 518 |
| *O. niloticus* | F | - | - | - | - | - | - |
| **RH2A2** | 53 (52) | 98 (97) | 123 (122) | 53 (52) | 98 (97) | 123 (122) | |
| *C. colonus* | F | T | Q | ? | ? | -13 | 505+/- tuning effect |

| | | | | | | | |
|---|---|---|---|---|---|---|---|
| *C. furcifer* | F | T | Q | ? | ? | -13 | 505+/- tuning effect |
| *C. fulva* | F | T | Q | ? | ? | -13 | 505+/- tuning effect |
| *O. niloticus* | T | S | E | - | - | - | - |
| **RH2B** | - | - | - | - | - | - | |
| *C. colonus* | - | - | - | - | - | - | 480 |
| *C. furcifer* | - | - | - | - | - | - | 480 |
| *C. fulva* | - | - | - | - | - | - | 480 |
| *P. amboinensis* | - | - | - | - | - | - | - |
| **LWS** | - | - | - | - | - | - | |
| *C. colonus* | - | - | - | - | - | - | 530 |
| *C. furcifer* | - | - | - | - | - | - | 530 |
| *C. fulva* | - | - | - | - | - | - | 530 |
| *R. guatemalensis* | - | - | - | - | - | - | - |

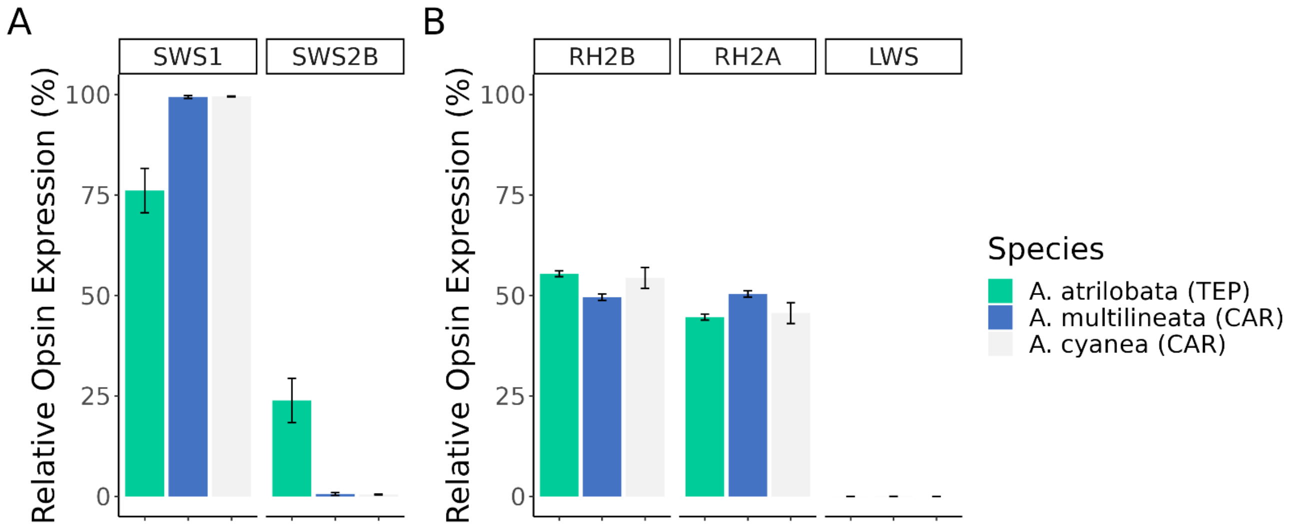


**Figure S12.** Damselfish relative expression levels of cone opsin genes expressed in (A) single cones; *SWS1* and *SWS2B,* where expression levels are given as percentages of the total single cone opsin expression and (B) double cones; *RH2A*, *RH2B* and *LWS,* where expression levels are given as percentages of the total double cone opsin expression. Error bars show +/- standard error.

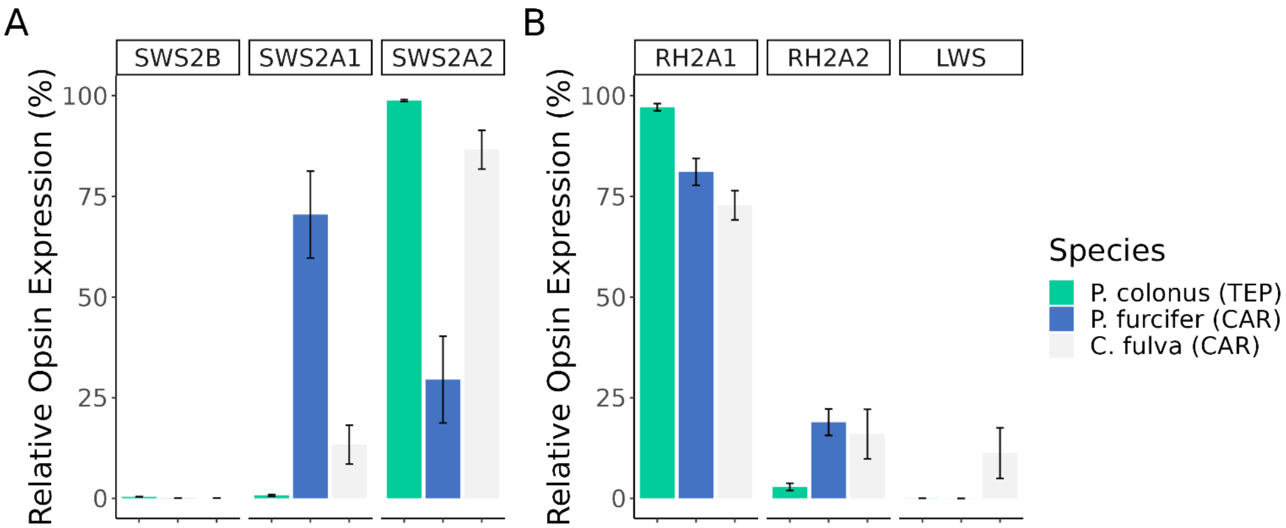


**Figure S13.** Grouper relative expression levels of cone opsin genes expressed in (A) single cones; *SWS1* and *SWS2B,* where expression levels are given as a percentage of the total single cone opsin expression and (B) double cones; *RH2A1*, *RH2A2* and *LWS,* where expression levels are given as percentages of the total double cone opsin expression. Error bars show +/- standard error.

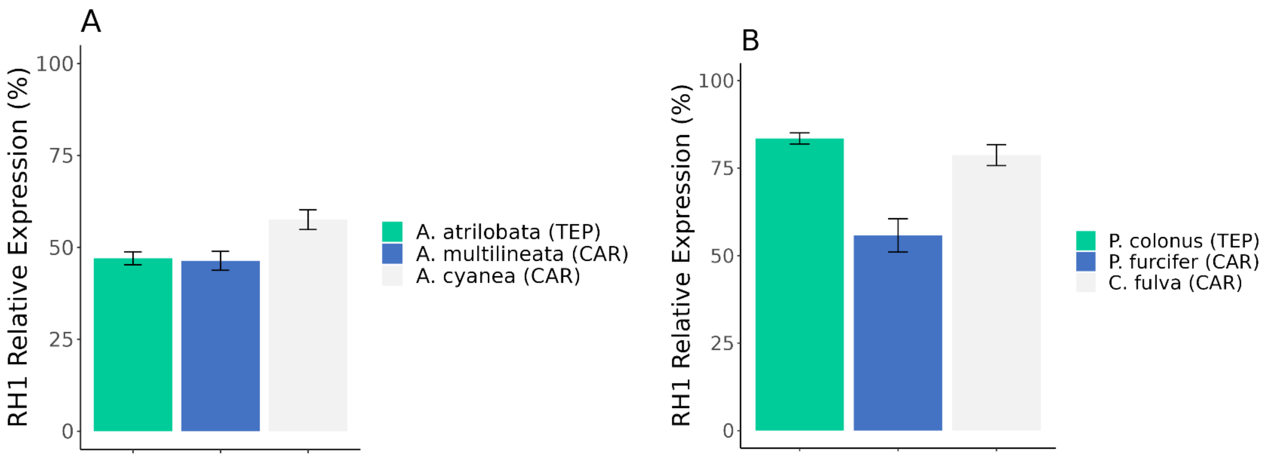


**Figure S14.** Relative expression levels of rod opsin *RH1* as a percentage of total visual opsin expression in the damselfish (A) and the grouper (B) geminate contrasts. Error bars show +/- standard error.

**Table S7.** Significantly differentially expressed genes between damselfish geminates in either phototransduction, circadian or non-visual opsin categories identified in the wider retinal transcriptome differential expression. The log2 fold change and Benjamini-Hochberg adjusted *p*-values from the DESeq2 differential expression analysis are reported. Rod and cone specific genes in the phototransduction pathway are highlighted blue or red respectively.

| Name | Function | baseMean | log2Fold Change | lfcSE | stat | *p*-value | padj |
|---|---|---|---|---|---|---|---|
| *GNAT1* | Photo-transduction | 366479.90 | 1.66 | 0.20 | 8.49 | 1.54E-09 | 1.13E-08 |
| *GNB1* | Photo-transduction | 104816.92 | 1.03 | 0.15 | 7.07 | 1.30E-04 | 5.83E-04 |
| *GNGT1* | Photo-transduction | 35253.79 | -6.50 | 0.27 | -24.12 | 4.11E-110 | 3.33E-108 |
| *CNGA1* | Photo-transduction | 5419.44 | 3.89 | 0.22 | 17.73 | 3.82E-54 | 7.75E-53 |
| *PDE6G* | Photo-transduction | 101709.54 | 3.63 | 0.16 | 22.10 | 3.22E-81 | 1.30E-79 |
| *GRK1* | Photo-transduction | 18306.06 | 2.66 | 0.21 | 12.41 | 3.78E-24 | 5.10E-23 |
| *GNGT2* | Photo-transduction | 78925.18 | 1.22 | 0.26 | 4.72 | 2.63E-03 | 7.61E-03 |
| *PDE6C* | Photo-transduction | 5758.70 | 3.31 | 0.40 | 8.37 | 5.81E-13 | 6.73E-12 |

| | | | | | | | |
|---|---|---|---|---|---|---|---|
| *PDE6H* | Photo-transduction | 126137.13 | 3.40 | 0.46 | 7.36 | 1.72E-10 | 1.74E-09 |
| *GRK7* | Photo-transduction | 3557.47 | 3.89 | 0.94 | 4.13 | 1.60E-04 | 6.81E-04 |
| *ARR3* | Photo-transduction | 120338.89 | 5.00 | 0.43 | 11.69 | 3.34E-26 | 5.41E-25 |
| *GUCA1A* | Photo-transduction | 67924.58 | -1.67 | 0.34 | -4.88 | 3.16E-04 | 1.28E-03 |
| *GUCA1B* | Photo-transduction | 11966.74 | 5.19 | 0.28 | 18.67 | 3.89E-64 | 1.05E-62 |
| *GUCY2D* | Photo-transduction | 3608.35 | 1.37 | 0.31 | 4.47 | 2.30E-03 | 7.20E-03 |
| *GUCY2F* | Photo-transduction | 3686.16 | 1.78 | 0.25 | 7.17 | 1.26E-07 | 7.30E-07 |
| *RGS9BP* | Photo-transduction | 358.56 | 2.08 | 0.52 | 4.02 | 1.14E-03 | 4.01E-03 |
| *GNB5A* | Photo-transduction | 1863.62 | -0.86 | 0.15 | -5.87 | 6.98E-03 | 1.82E-02 |
| *CLOCK A* | Circadian | 400.51 | 1.57 | 0.36 | 4.34 | 1.56E-03 | 5.26E-03 |
| *PER1* | Circadian | 170.05 | 2.44 | 0.61 | 4.02 | 6.85E-04 | 2.64E-03 |
| *PER2* | Circadian | 97.87 | 3.29 | 0.51 | 6.47 | 2.01E-08 | 1.25E-07 |
| *PER3* | Circadian | 13.96 | 8.25 | 1.25 | 6.59 | 3.00E-10 | 2.70E-09 |
| *CRY1A* | Circadian | 59.87 | 1.95 | 0.38 | 5.11 | 7.25E-05 | 3.45E-04 |

| | | | | | | | |
|---|---|---|---|---|---|---|---|
| *CRY1B* | Circadian | 61.04 | 2.96 | 0.42 | 7.14 | 1.49E-09 | 1.13E-08 |
| *CRY5* | Circadian | 15.11 | -1.84 | 0.57 | -3.23 | 9.40E-03 | 2.38E-02 |
| *RORB* | Circadian | 2743.39 | -1.39 | 0.35 | -3.95 | 5.62E-03 | 1.57E-02 |
| *CIART* | Circadian | 17.79 | 6.64 | 1.23 | 5.39 | 3.19E-07 | 1.72E-06 |
| *CSNK1DB* | Circadian | 607.74 | 1.13 | 0.26 | 2.47 | 1.35E-02 | 3.64E-02 |
| *NOCTA* | Circadian | 1179.70 | -1.52 | 0.36 | -4.21 | 2.32E-03 | 7.20E-03 |
| *RGRA* | Non-visual Opsin | 1102.21 | 1.82 | 0.42 | 4.30 | 9.00E-04 | 3.31E-03 |
| *OPN4B* | Non-visual Opsin | 59.08 | -1.81 | 0.47 | -3.89 | 2.40E-03 | 7.20E-03 |
| *VAL-OPSIN-A* | Non-visual Opsin | 28.00 | -2.82 | 0.39 | -7.16 | 1.93E-09 | 1.30E-08 |
| *OPN6A* | Non-visual Opsin | 1213.15 | 2.17 | 0.35 | 6.21 | 8.72E-07 | 4.42E-06 |

**Table S8.** Significantly differentially expressed genes between grouper geminates in either phototransduction, circadian or non-visual opsin categories identified in the wider retinal transcriptome differential expression. The log2 fold change and Benjamini-Hochberg adjusted *p*-values from the DESeq2 differential expression analysis are reported. Rod and cone specific genes in the phototransduction pathway are highlighted blue or red respectively.

| Name | Function | baseMean | log2Fold Change | lfcSE | stat | *p*-value | padj |
|---|---|---|---|---|---|---|---|
| *GNAT2* | Photo-transduction | 54307.29 | 2.01 | 0.35 | 5.67 | 1.02E-05 | 2.67E-04 |
| *GUCA1C* | Photo-transduction | 2759.55 | 3.68 | 0.98 | 3.75 | 6.09E-04 | 6.25E-03 |
| *GNB5A* | Photo-transduction | 1868.75 | -1.43 | 0.25 | -5.64 | 1.22E-04 | 1.43E-03 |
| *TIMELESS* | Circadian | 31.59 | 3.32 | 0.68 | 4.85 | 1.88E-05 | 3.08E-04 |
| *NOCTA* | Circadian | 1861.03 | 2.38 | 0.49 | 4.82 | 7.09E-05 | 9.69E-04 |
| *NPAS2* | Circadian | 1360.66 | 2.36 | 0.64 | 3.68 | 1.86E-03 | 1.70E-02 |
| *RGRB* | Non-visual Opsin | 3335.93 | -3.39 | 0.63 | -5.40 | 2.09E-06 | 8.57E-05 |
| *RRH* | Non-visual Opsin | 389.07 | -4.00 | 0.73 | -5.45 | 9.35E-07 | 7.66E-05 |
| *OPN4A* | Non-visual Opsin | 164.17 | 3.39 | 0.69 | 4.93 | 1.30E-05 | 2.67E-04 |

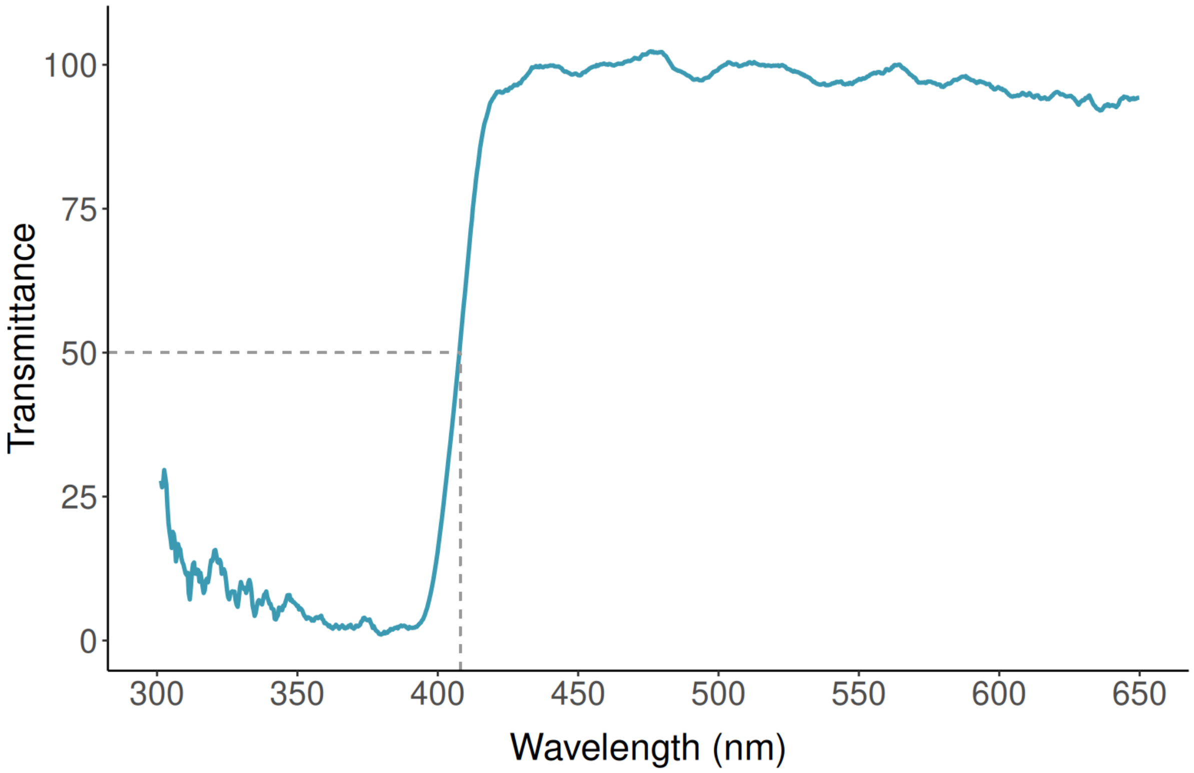


**Figure S15.** Lens transmittance in *Cephalopholis furcifer:* the wavelength at which 50% of the maximal transmittance (T50) is reached is indicated. Note that very little light is transmitted below 400nm (i.e. ultraviolet region).

### *Transcriptome Quality*

We assessed the extent to which each of the transcriptome assemblies recovered single-copy orthologs present in the Actinopterygii database ($n$ = 3640) (Fig. S16). Since here we were reconstructing the transcriptomes of a single, highly specific tissue, we did not expect to recover the full complement of single-copy orthologs, however we did expect consistency across the transcriptome BUSCO scores for each species. Across the six transcriptome assemblies, the number of complete single-copy orthologs ranged from 76.1 - 85.2%, fragmented from 2.9 - 4.7% and missing 11.9 - 19.4%. The number of complete BUSCO's recovered was relatively consistent across all six species and they were also comparable with de novo transcriptome assemblies of other coral reef fishes even where these have been based on more general tissues (Bernal et al., 2020; Maytin et al., 2018).

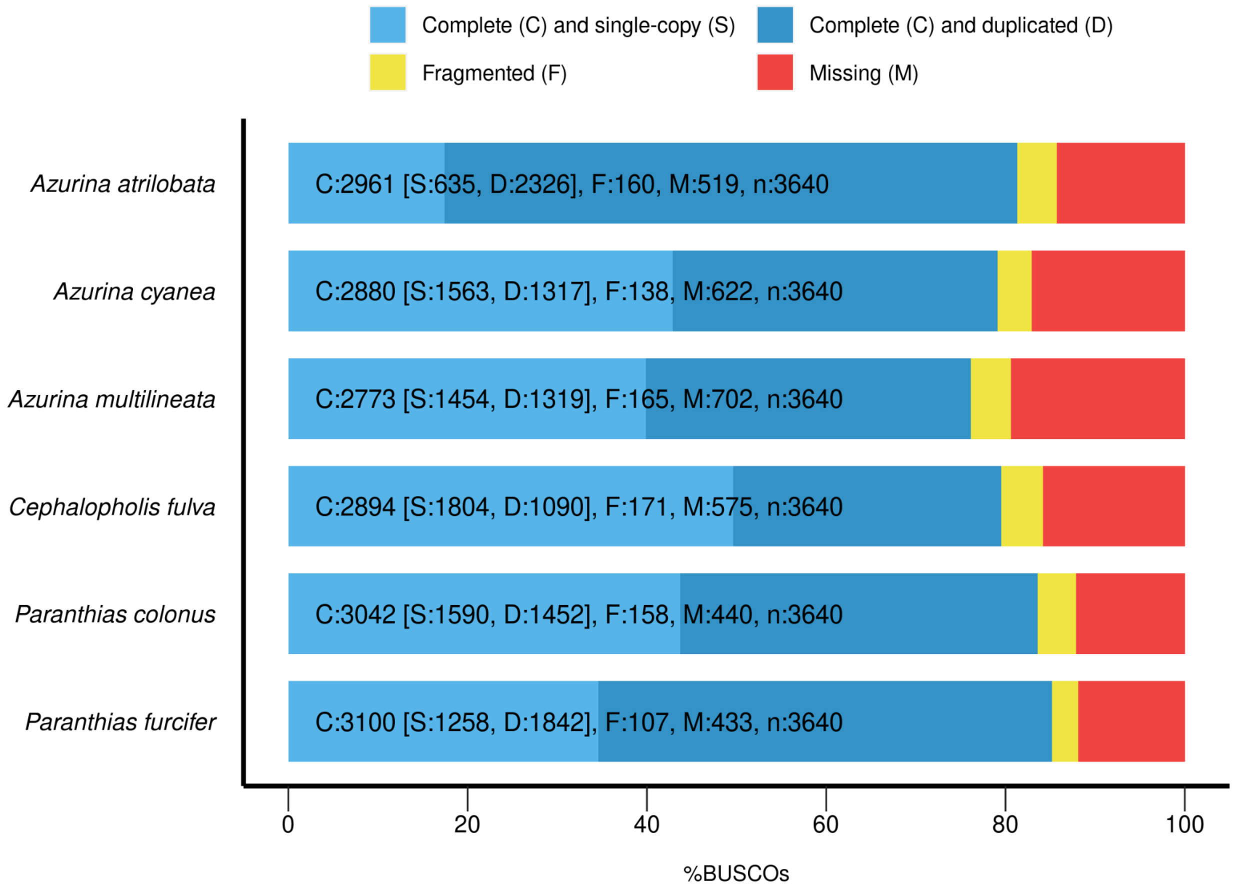


**Figure S16.** BUSCO assessment results for each species' transcriptome assembly, benchmarked against the Actinopterygii single-copy ortholog database (odb9).

For each assembly we calculated basic contig metrics which are summarised in Supplementary Table S10. These showed high consistency across the assemblies. To assess the structure of the transcriptome assemblies and distribution of contig lengths we used ExN50 curves which show the percentage of the normalised expression data against the N50 values. The peaks and corresponding contig lengths were consistent across all the species transcriptomes (Fig. S17). The further shifted to the right the ExN50 peak, the better coverage, and peaks towards ~80-90% expression are indicative of a very high quality assembly where deeper sequencing is unlikely to further improve assembly quality. The consistent peaking of all the ExN50 plots at a reasonably high expression percentile suggested there was not a high proportion of transcripts with a low rate of expression present in our data. This indicates the sequencing depth here was adequate and we would not expect deeper sequencing to markedly improve the assembly quality.

**Table S9.** Basic contig metrics calculated for all six species *de novo* transcriptome assemblies. Contig N50 is a measure of assembly contiguity where the N50 value represents the smallest contig length in which 50% of assembled transcript nucleotides are found.

| Species | *A. atrilobata* | *A. multilineata* | *A. cyanea* | *C. colonus* | *C. furcifer* | *C. fulva* |
|---|---|---|---|---|---|---|
| Total Number of Assembled Bases | 444789548 | 354442414 | 355435673 | 499145510 | 334605358 | 327444762 |
| Total Number of Contigs | 385737 | 295709 | 287630 | 478677 | 283131 | 276504 |
| Contig N50 | 2338 | 2446 | 2604 | 2105 | 2592 | 1496 |
| Mean Contig Length | 1153 | 1199 | 1236 | 1043 | 1182 | 1184 |

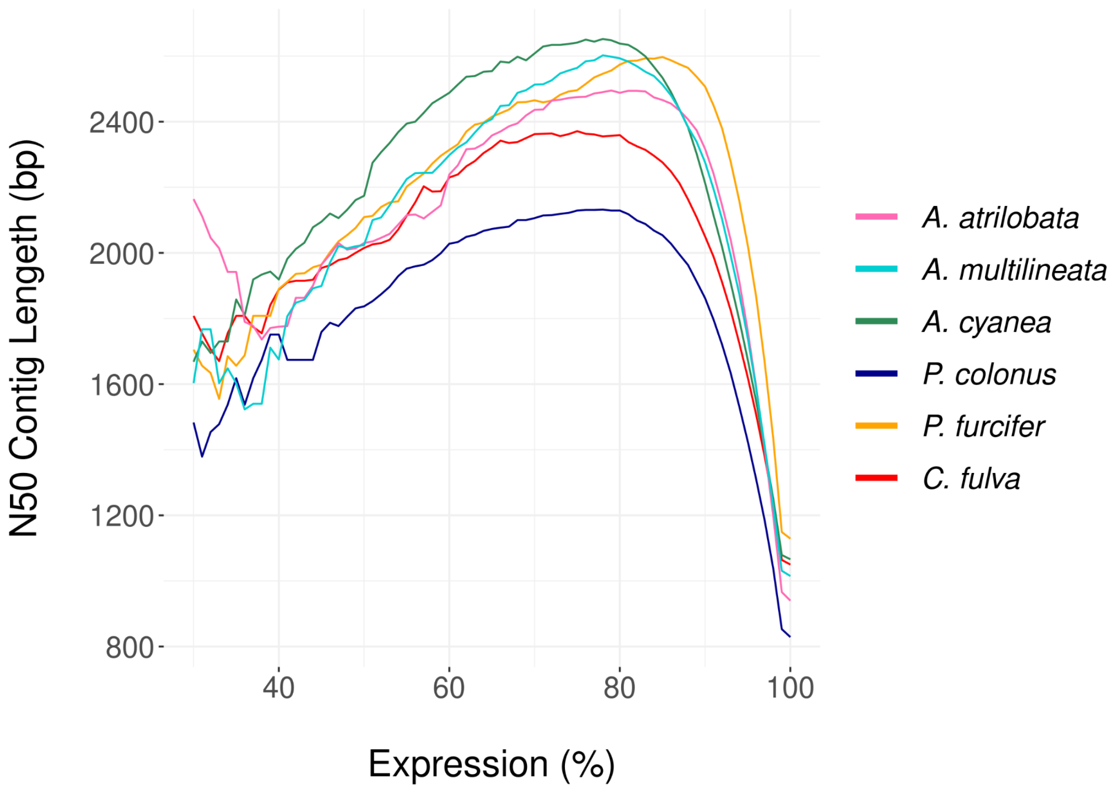


**Figure S17.** ExN50 curves of all six transcriptome assemblies. N50 contig length values (the smallest contig length in which you find 50% of assembled transcript nucleotides) but limited to the top x% of the normalized expression data. ExN50 curves show similar distributions, peaking between 75-85% expression which corresponds to a N50 contig length of over 2000bp in all assemblies. Peaks at higher x% are indicative of sufficient sequencing depth and higher quality assembly.
